# Machine Learning to Predict Continuous Protein Properties from Simple Binary Sorting and Deep Sequencing Data

**DOI:** 10.1101/2023.06.09.544229

**Authors:** Marshall Case, Matthew Smith, Jordan Vinh, Greg Thurber

## Abstract

Proteins are a diverse class of biomolecules responsible for wide-ranging cellular functions, from catalyzing reactions and recognizing pathogens to forming dynamic cellular structure. The ability to evolve proteins rapidly and inexpensively towards improved properties is a common objective for protein engineers. Powerful high-throughput methods like fluorescent activated cell sorting (FACS) and next-generation sequencing (NGS) have dramatically improved directed evolution experiments. However, it is unclear how to best leverage this data to characterize protein fitness landscapes more completely and identify lead candidates. In this work, we develop a simple yet powerful framework to improve protein optimization by predicting continuous protein properties from simple directed evolution experiments using interpretable machine learning. Evaluated across five diverse protein engineering tasks, continuous properties are consistently predicted from readily available deep sequencing data. To prospectively test the utility of this approach, we generated a library of stapled peptides and applied the framework to predict and optimize both affinity and specificity. We coupled integer linear programming with the interpretable machine learning model coefficients to identify new variants from experimentally unseen sequence space that have desired properties. This approach represents a versatile tool for improved analysis and identification of protein variants across many domains of protein engineering.

## Introduction

A longstanding goal of biochemistry has been to map the sequence of a protein to its structure and function.^1^ However, the complex biophysics that govern the protein fitness landscape, including how a protein folds and how its structure influences function, make the coupling of sequence to function an extremely difficult task. Protein engineers thus often focus on a much smaller subdomain of the protein fitness landscape, using the confined resources of experimental protein science to explore variants close to a known functional protein with the goal of incrementally improving function. A common and extremely powerful approach is directed evolution, where a protein is encoded by DNA, expressed by cells, and assayed by magnetic or fluorescent activated cell sorting (MACS or FACS) and, more recently, next generation sequencing (NGS) to identify variants with improved fitness. While these techniques represent powerful tools in the protein engineering arsenal, it is unclear how to best leverage information from deep sequencing towards the optimization of protein variants. A method capable of generating both fitness estimates from directed evolution experiments and predictions of sequences with higher activity would greatly expand the power and efficiency of directed evolution experiments.

The combination of directed evolution and next generation sequencing (NGS) has enabled protein engineers to rapidly evaluate millions to billions of protein variants in a highly focused manner. With maintenance of the genotype-phenotype connection, any technique that manipulates DNA in a high-throughput manner can be applied to design focused protein variant libraries and assay protein function.^2, 3^ Techniques like mRNA display and phage display can evaluate the largest libraries, although their small size precludes them from sorting approaches such as FACS.^4^ Cell surface display techniques, which use bacteria or yeast, enable facile measurement of the interaction between protein variants with soluble proteins which can be used for assaying binding affinity in high-throughput sorting and sequencing technologies.^5^ Coupling FACS with cell surface display technologies allows for the selection of rare protein variants among a large library with extreme selectivity.^6, 7^ These techniques have enabled a wide range of protein engineering campaigns, from affinity maturation of protein-protein interactions to highly enantioselective enzymes.^8, 9^ However, one challenge with these large libraries is how to identify the best lead molecules from the hundreds to thousands of observed sequences in the final sorted population. Traditional approaches for lead molecule identification select variants according to their abundance in the enriched library under the assumption that higher enrichment is indicative of higher function.^10–12^ One downside to this approach is that optimal rare variants are excluded from selection and more complex descriptions of how mutations contribute to protein function are difficult to ascertain.^11^ Application of NGS to the output pool of a protein variant sort improves the accuracy of clone frequency, but frequency rarely correlates with protein properties directly.^13–, 15^ These challenges arise from sources of error that are difficult to eliminate: variation in cell-to-cell growth, PCR/cloning biases, sequencing errors, and FACS instrument noise.^16, 17^ With additional sequencing of the input library, enrichment ratios can be calculated, which improves the accuracy of protein property prediction.^18, 19^ Despite these improvements, there is still little consensus on the best experimental design and analysis of these directed evolution experiments.

Several approaches have been proposed to mitigate these sources of error and enable the prediction of quantitative protein properties from high-throughput sorting experiments. Deep mutational scanning (DMS) measures the enrichment of many variants. However, several challenges exist; their accuracy in resolving affinity is often limited to a narrow linear region (∼10X dynamic range), the results are sensitive to the sorting conditions, stability, and expression effects, and the outcomes can differ from true quantitative measurements of binding affinity (equilibrium dissociation constants or K_D_’s).^20, 21^ Sort-seq aims to address noise from sorting by using multiple bins across the entire fluorescent channel, followed by deep sequencing, to infer the distribution of each sequence in fluorescent space.^22^ These techniques, while often successful, require more sorting time and 8-12 fold increased deep sequencing throughput and still have a narrow range of resolution. Several more sophisticated sorting techniques address these issues: SORTCERY creates a rank ordering of affinities by sorting cells according to their binding and expression at a single concentration;^23^ amped SORTCERY further improves this technique by converting rank order to free energy changes by adding titration standards;^24^ TiteSeq sorts protein variants at multiple ligand concentrations and fits the affinity to the fraction bound.^21^ These methods leverage additional sorting and sequencing to improve the predicted outcomes. In this work, we seek to utilize deep sequencing with interpretable machine learning approaches to determine if we can predict continuous protein properties (like affinity) from binary sorting data (positive versus negative sorting).

## Methods

### Curation of NGS Data for Validation

Five datasets were used to test the simple method of using binary labels to predict continuous properties. The datasets and brief descriptions are given below.

#### 1. Adams et al. 2016^21^

NGS data was downloaded from their GitHub repository: https://github.com/jbkinney/16_titeseq. The read counts and CDR^1H^ and CDR^3H^ sequences for each clone were extracted and aligned using in-house python scripts. Read counts were converted to frequencies.

#### 2. Starr et al. 2022 and Greaney et al. 2021^25, 26^

NGS data was downloaded from their GitHub repository: https://github.com/jbloomlab/SARS-CoV-2-RBD_DMS_variants. The data for the Delta mutation is stored in a different repository: https://github.com/jbloomlab/SARS-CoV-2-RBD_Delta. Due to limits in Illumina paired end reading length, each sequence was given a unique molecular barcode, which was sequenced in high depth, but each full-length sequence was sequenced with its unique barcode separately. The sequences and their TiteSeq profiles were associated with their corresponding barcodes and read counts were converted to frequencies. In the current method, sequences with more than one mutation were not discarded.

#### 3. Makowski et al. 2022^27^

Processed data was downloaded from their GitHub repository: https://github.com/Tessier-Lab-UMich/Emi_Pareto_Opt_ML. Raw data was available from their repository.

#### 4. Sarkisyan et al. 2016^28^

Like Starr et al. 2022, the GFP sequence is too long for high-depth Illumina sequencing, and therefore the authors gave each sequence a unique molecular barcode. We downloaded the accurate full length protein sequences, their matching unique barcodes, and the high-depth sequencing of Sort-seq data from their repository: https://figshare.com/articles/dataset/Local_fitness_landscape_of_the_green_fluorescent_ protein/3102154. The read accuracy on the barcodes was low and the authors used a Levenshtein distance of <= 1 to connect barcodes that were close but not identical to the full protein sequence. We used the Levenshtein module with in-house python scripts to cluster sequences to their barcodes, which were available at https://pypi.org/project/python-Levenshtein/. After clustering, sequences and their barcodes were merged with their Sort-Seq distributions like Starr et al. Read counts were converted to frequencies.

#### 5. Jenson et al. 2018^24^

NGS data was obtained from their GitHub repository: https://github.com/KeatingLab/sortcery_design. The peptides’ short lengths permitted high depth deep sequencing and thus counts were directly converted to frequencies without further preprocessing.

### Binarization of FACS/NGS Data

The variety of factors that influence the design of an experiment makes it challenging to generalize a sorting and sequencing workflow for any given protein engineering campaign. Each of these projects were analyzed by a different group, using different cell sorters, expression platforms, sequencing instruments, and protein types among other parameters (see Table S1 for dataset property summaries). Thus, controlling each of those parameters in our data processing workflow was an important consideration towards the application of this approach to existing datasets and new targets alike. Many of the experiments use sophisticated sorting techniques to infer quantitative protein properties. We simulated a simple binary sort experiment by truncating the dataset such that it only includes the top or bottom 20% of sorted sequences (or as close as possible). This subsample of sequencing data approximates a simple sorting campaign from these quantitative sorting techniques. For example, **Sarkisyan et al.** contains sequencing data of GFP variants that were sorted into 8 bins; to simulate a simple binary sort, we aggregated the top two bins as positive and the bottom two bins as negative. For TiteSeq experiments (Starr 2022, Greaney 2021, and Adams 2016) we only included data from sorts that used ligand concentrations near the average K_D_ of the library (10^-9^, 10^-9^, and 10^-8^ M respectively). Because the K_D_ of a library can be readily obtained from low-throughput flow cytometry experiments, sorting at the K_D_ of the library is a feasible approach to yield the largest separation between high and low affinity variants.^17^ This was 10^-8^ M for the COVID datasets, this was 10^-8^ M and 10^-9^ M, respectively. For the **Makowski dataset**, data was provided as a positive and negative dataset with varying cutoffs for each selection.

### Machine Learning Method

In all cases, in-house python scripts were used to perform the data preparation and modeling on each of the datasets. Scikit-learn (https://scikit-learn.org/stable/) was used for linear discriminant analysis (LDA), one-hot encoding, scaling label vectors, and other pre-processing steps. Pandas (https://pandas.pydata.org/) and NumPy (https://numpy.org/) were used to handle sequencing and numerical data. PyTorch (https://pytorch.org/) was used to train neural network models.

First, sequences were one-hot encoded, eliminating positions that were not randomized in the study or appeared with very low abundance. Then, we calculated the frequencies of each sequence for the high- and low- protein property, and a multi-sequence alignment (MSA) was performed to ensure every vector was the same length and columns corresponded to the correct residues. The data was split into positive and negative groups by computing the ratio of high- and low- frequency of each clone and selecting a percentile cutoff. Initial percentiles were chosen as the top or bottom 20% of sequences, setting any sequences that contained zero frequency in the low property pool to the maximum ratio observed and any sequences that contained zero frequency in the high property pool to the minimum ratio observed. Positive (‘1’) and negative (‘0’) labels were assigned accordingly. The one-hot encoded protein sequences and their labels were then split into an 80:20 training:test split. The test set was held aside until all analyses were complete and used to validate the model training process. In later analyses, to explore the hyperparameter space of these cutoff parameters, we tested all combinations of the read count, replicate count, and ratio percentile and measured the change in modeling performance. Sensitivity to training:test splitting and the ratio of positive negative labels was tested by performing five-fold cross validation using SciKit Learn’s ShuffleSplit function.

We selected linear discriminant analysis for several reasons. First, this method has previously been shown to predict continuous properties from binary sorting data.^27^ Next, hyperparameter optimization for this model was straightforward, as the Sci-Kit Learn implementation of LDA has very few parameters, including the solver (‘svd’ was the only one to converge consistently), n_components (which is fixed to 1 for projection to a single dimension to correlate with protein properties), and tol (which did not change the outcome). Another benefit of using LDA is its simplicity; the linear nature of the model allows for the direct interpretation of how certain residues contribute to function. While we also evaluated several other models that can create an internal continuous representation for classification (such as support vector classifiers, with the option of using different kernels), we found that LDA models trained much faster. The transform method was used to project data into the 1-dimensional LDA projection after training. Because LDA is a classification model and does not have a regression analog, we used ridge regression, a modified version of linear regression that penalizes large weights, to compare LDA projections to models trained on continuous data. Furthermore, ridge regression did not result in extreme overfitting that was observed by regular linear regression (data not shown). Finally, ridge regression has been shown to be a powerful modeling technique for protein engineering tasks.^29^

Neural network models were used to evaluate whether non-linear models would capture additional useful information that linear models are unable of modeling, as proposed previously.^27^ Standard fully connected, feed forward networks were used with dropout p = 0.5 as shown to be effective in the literature.^30^ The hidden size (32-256) and number of layers (1-3) did not dramatically affect the results and we ultimately chose the midpoint for both, 128 and 2 respectively (data not shown). We used 700 epochs and a batch size of 32 was for all datasets. Binary Cross Categorical Entropy Loss was used as the loss function, and Stochastic Gradient Descent optimizer with a learning rate of 0.01 was used for all datasets. Training was done on a Nvidia Tesla V100 and typically took between 5 minutes and 2 hours depending on the size and complexity of the dataset.

### Stapled peptide cell sorting, sequencing, and flow cytometry

Experimental stapled peptide libraries targeting B cell lymphoma 2 (Bcl-2) proteins were used to evaluate the computational methods on novel datasets. These libraries were sorted and sequenced as described previously. (Case 2023, manuscript in progress) In brief, a computational library of BIM mutants (a non-specific anti-apoptotic peptide) was designed and transformed into bacteria that displays stapled peptides (see Table S2 for mutagenesis codons, Table S3 for sampled amino acids, and Table S4 for library primers).^31, 32^ This library was sorted using a combination of MACS and FACS as follows: one round of expression MACS, two rounds of affinity MACS, two rounds of affinity FACS, and two rounds of specificity FACS. Two of such libraries were sorted in parallel: one towards Bcl-x_L_ and another towards Mcl-1. These libraries were deep sequenced using Illumina NovaSeq S4, demultiplexed, merged using NGMerge, and analyzed using in-house python scripts (see Table S5 for NGS primers).^33^ Each peptide sequence was identified by aligning the DNA with the scaffold eCPX protein and then translating the peptides in the corresponding open reading frame. Peptide sequences and their frequencies were aligned across all rounds of sorting, and sequences that had mutations not specified by the original library design were removed (∼10% of all sequences). Sequences from the four rounds of FACS were denoted as ‘hits’ and sequences from the expression sort were denoted as ‘not hits’ (see Table S1 for dataset summary). Then, the ratio of each round of FACS to the expression was computed and fed into the machine learning pipeline. LDA models were trained identically to the other datasets.

A smaller number of peptide sequences were expressed on the surface of bacteria and measured in low-throughput flow cytometry experiments. To evaluate whether LDA projections were predictive of continuous properties, we expressed 57 stapled peptides on the surface of bacteria from various rounds of sorting (Mcl-1 FACS 2, 3, or 4, and Bcl-x_L_ FACS 2 or 4) to capture a wider distribution of specificities: peptides from later in the rounds of sorting should have more specificity while those from earlier rounds should be less specific if sorting enriched towards higher performing sequences. We then measured their binding at the approximate K_D_ of the wild type sequence in triplicate (1nM and 10nM for Mcl-1 and Bcl-x_L_ respectively). Fraction bound was calculated by normalizing to expression and dividing by a saturated binder (BIM-p5 at 250nM).^32^ LDA projections were calculated and compared to continuous values identically to the other datasets.

### SORTCERY

To get continuous estimates of binding properties from cell sorting, peptides from the final round of FACS for both Mcl-1 and Bcl-x_L_ were evaluated using SORTCERY. Peptides were incubated with either Mcl-1 or Bcl-x_L_ at 1nM and sorted into twelve bins following the protocol from Reich et al.^23^ Briefly, cells labeled with target Bcl-2 protein and anti-HA display tag were sorted into twelve bins along the ‘axis of affinity’, the diagonal gates that resolves the fraction bound. To compare the SORTCERY value with those measured from binary sorting, we computed the gate score of each sequence as described in the original work.^23^ Each of these gates were collected individually and processed for deep sequencing as described previously. The deep sequencing data from these experiments was processed identically to the stapled peptide libraries as above.

### Sequence Optimization via Integer Linear Programming

To optimize protein sequences, we applied integer linear programming (ILP), an approach that solves an objective problem given discrete input variables and constraints. Compared to other techniques that maximize an objective given an input, ILP scales more efficiently with a large number of samples and does not rely on iterative predict and test loops that require additional experimental resources.^34–36^ Furthermore, ILP is directly amenable to multi-objective optimization through the addition of inequality requirements.^24^ We set up this problem using the PuLP python module.^37^ First, we defined the objective as maximizing the dot product of the model coefficient vector and the positions and amino acid constraints as defined by the library design. This objective is the maximization of the confidence of binding for a given sequence. Next, we constrained the optimization by only allowing one amino acid at each position, requiring that each peptide had two azidohomoalanine residues (responsible for peptide stapling), and that the two stapled residues were at a distance as specified by the library design (i,i+7). Finally, we formulated the problem as a multi-objective problem by adding the additional constraint that the dot product of the off-target coefficients and peptide sequence was in the non-binding regime.

## Results

### Overview of Method

Despite significant efforts to gather quantitative data from high throughput sorting, most directed evolution campaigns rely on basic metrics of protein fitness. We utilized a simple workflow to extract continuous protein properties from NGS datasets while keeping the experimental design simple and affordable **(Figure 1)**. To accomplish this task, we generated binary labels from enrichment ratios, trained machine learning models using these binary labels to infer continuous protein properties,^27^ and optimized protein sequence and function beyond experimentally sampled space into unseen sequence space.^24^ We hypothesized that continuous protein properties can be obtained from simple sorting and sequencing analyses for three primary reasons. First, because cell sorting is a stochastic process, cells sorted into discrete bins are sampled from an underlying continuous distribution. Thus, cells sorted in a binary manner may allow inference of this distribution.^38^ Second, biased sampling towards the most and least functional variants may allow models to ‘interpolate’ function of intermediate fitness. Finally, sampling many epistatically interacting motifs may allow inference between them.^39^ We also hypothesized this approach would work across multiple protein engineering objectives, including affinity maturation, fluorescence, deep mutational scanning, and specificity.

**Figure 1:**
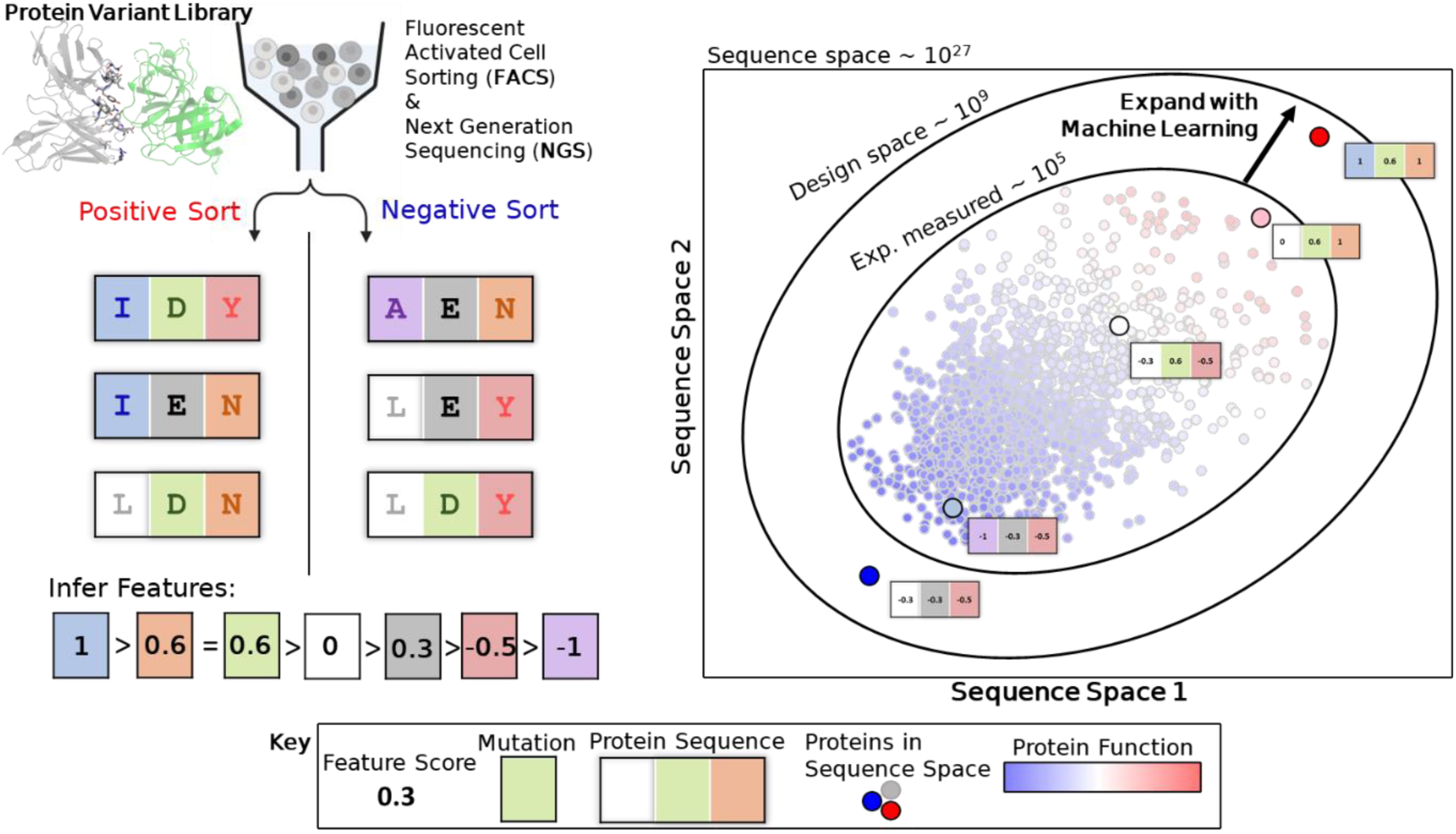
Overview of the protein engineering workflow. A library of protein variants is expressed on the surface of cells and sorted according to its fluorescent property (binding fluorescent labeled molecules, intrinsic fluorescence, etc.). Sorted cells are deep sequenced, and using machine learning, continuous scores for each mutation and the overall sequence are calculated. Finally, these data are used to inform protein fitness landscapes and optimization of protein properties in engineering efforts.

To validate the approach, we aggregated data from multiple protein engineering campaigns that fulfilled two criteria: 1. they had many data points of multi-mutant proteins from a sorting campaign and 2. they had measured many continuous protein properties among these variants. These datasets were the fitness landscape of GFP,^28^ the directed evolution of a fluorescein-binding scFv,^21^ and the fitness landscape of SARS-COV-2 Spike protein .^25, 26^ Because the co-optimization of multiple properties is often needed, we also gathered datasets that design high-affinity and high- specificity monoclonal antibodies^27^ and highly specific peptides between three B cell lymphoma 2 (Bcl-2) proteins.^24^

### Data processing pipeline for varying protein variant libraries and sorting schemes

The modular data processing and machine learning pipeline to analyze multiple protein variant libraries consists of multiple steps **(Figure 2)**. First, a library of protein variants is sorted, and the ratio of the positive to negative gate frequencies is calculated for all sequences based on the deep sequencing data. If a sequence found in the positive gate but was unobserved in the negative gate, the ratio was set to the maximum observed; conversely, if a sequence found in the negative gate was unobserved in the positive gate, the ratio was set to 0. We hypothesized this ratio scheme balances the information gained from enrichment ratios while still including clones that were overwhelmingly enriched or depleted. Labels (‘1’ for high performing variants and ‘0’ for low performing variants) were assigned by determining a cutoff based on the average ratio (percentile ≥0.8 and ≤0.2 respectively) across how many replicates they appeared in (≥2). We hypothesized that while splitting the positive and negative labels at the 50^th^ percentile would increase the data size, sorting noise around the midpoint would confound information gained from binary ratios (Figures S1-5). We also hypothesized that removing sequences with 1 replicate would further reduce noise from sorting. Initial estimates of these parameters were chosen to balance the size of the dataset, the strictness of inclusion, and the confidence of the sequencing data. Having easily modifiable parameters for label assignment serves as both a tool for sequencing quality processing and a powerful hyperparameter in the subsequent machine learning steps (see Figures S1-5 for hyperparameter effect on dataset size).

**Figure 2:**
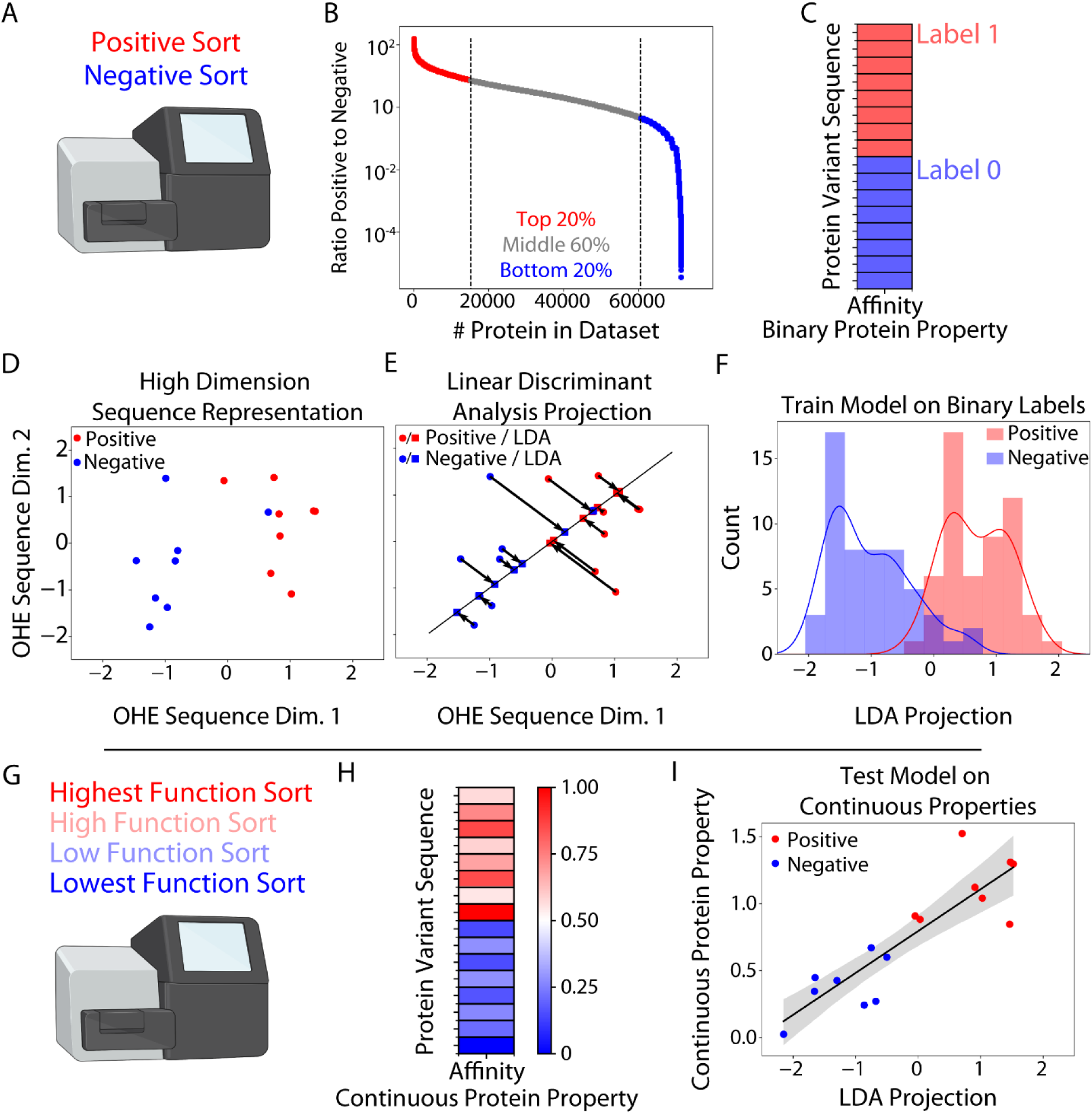
Deep sequencing, data pre-processing, and machine learning overview. The most and least functional protein variants from the binary sort are sequenced **(A)** and the ratio of sequence reads in the positive versus negative gate is calculated **(B)**. Binary labels are assigned to each sequence according to its ratio **(C)**; the label thresholds are easily modified depending on the library construction, sorting strategy, and sequencing data quality. Protein sequences are one hot encoded for machine interpretability **(D)** before being used to train a Linear Discriminant Analysis (LDA) model **(E)**, which is evaluated on a hold-out test set **(F)**. Then to calibrate the LDA model, continuous protein properties are obtained either from a quantitative sort (SORTCERY, Sort-Seq, or TiteSeq) or from low throughput measurements (flow cytometry titrations, ELISA, etc.) **(G,H)**. Finally, the projections from the LDA model are used to predict continuous protein properties **(I)**.

Armed with a dataset of sequences and binary function labels, a linear discriminant analysis (LDA) machine learning model was trained because it fulfilled two criteria: it could perform classification of sequence with its function label, and it had an internal continuous measurement that could be used to correlate with continuous properties. Because LDA models project high dimensional sequence data to maximize class separation, the final projection is a continuous representation that has been previously shown to correlate with continuous properties.^27^ The model was trained and tested by splitting the sequencing data into train and test sets randomly (80:20 train:test). To evaluate whether the weights learned by the LDA model correlated to meaningful continuous properties, a subset of the sequences were assayed for their property from a lower throughput but more accurate technique. For all but the Makowski dataset, this was a quantitative cell sorting experiment, and otherwise a low throughput measurement of affinity or specificity via flow cytometry with individual sequences. We then predicted the continuous properties of proteins by comparing the projections from LDA with actual continuous measurements.

### Binary labels predict protein properties with equal correlation power

To evaluate whether the LDA models trained on binary sorting data inferred meaningful features of the protein properties, we curated five datasets as described in the methods (see Table S1 for dataset summaries). Using data from each of the these, we compared the measured continuous properties of protein variants to their predicted values from LDA models trained on binary sorting data, as shown in **Figure 3A** (left) for the **Sarkisyan et al.** dataset. We next sought to determine the performance of a comparable model trained on continuous data. Continuous data is more expensive and/or complicated to obtain but presumably is more information rich. Therefore, we hypothesized models trained on continuous data would have stronger correlative power. To evaluate this hypothesis, we trained Ridge regression models, which have been previously shown to be powerful linear models that are not prone to over-fitting.^29^ We then compared the ability for both LDA and Ridge models to predict continuous properties (**Figure 3A**, right). Surprisingly, for the **Sarkisyan 2016 dataset**, the LDA models performed similarly to the Ridge regression models as evidenced by a similar Spearman’s ρ (0.846 for the LDA model and 0.855 for the Ridge regression model).

**Figure 3:**
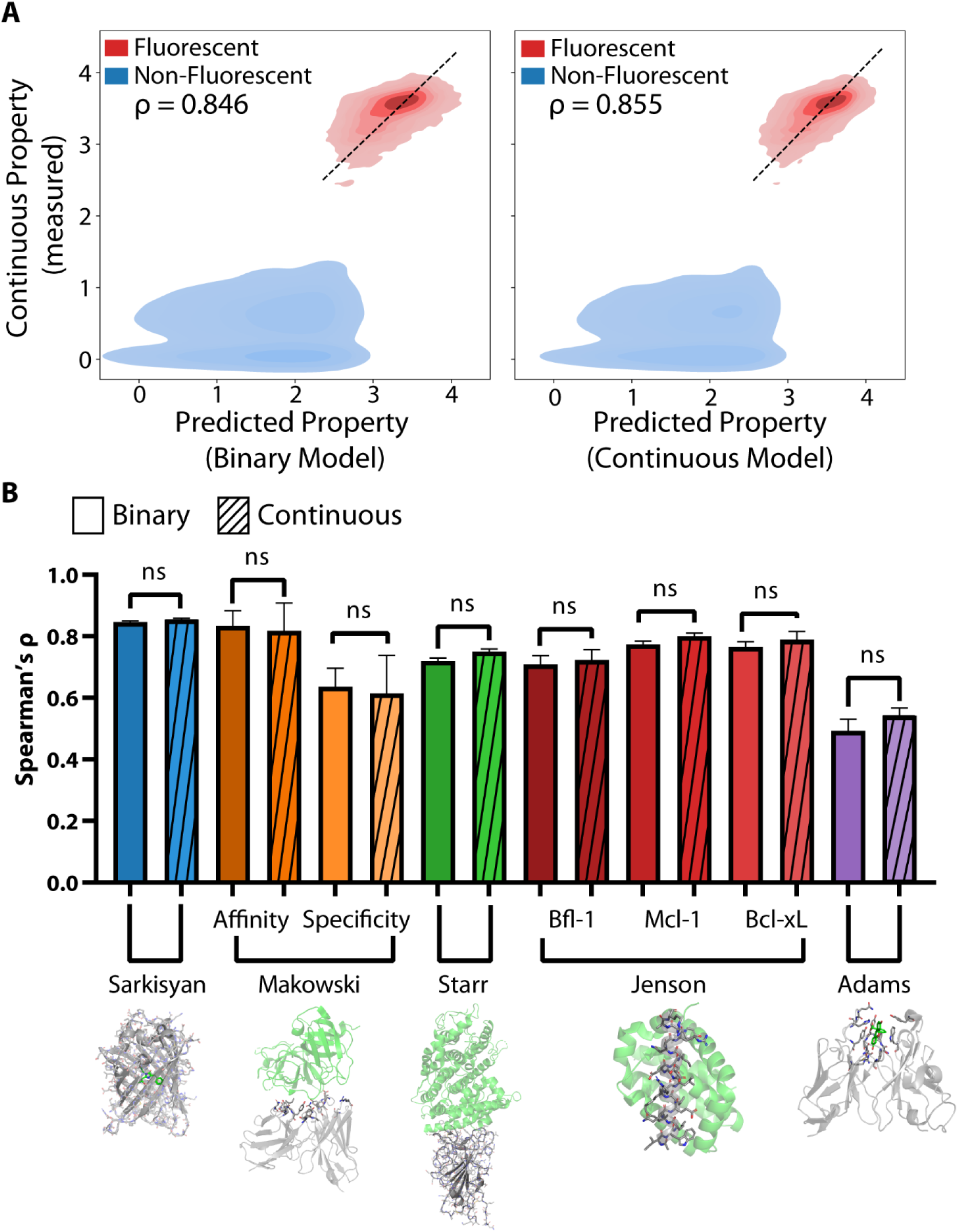
Predictions from models trained on binary data are highly correlated with continuous protein properties and equally powerful as models trained on continuous data. Evaluated on the Sarkisyan data, LDA models trained on binary data **(A, left)** or Ridge models trained on continuous data **(A, right)** are correlated with fluorescence. Across five protein engineering datasets, models trained on binary data are equally predictive of continuous properties **(B).**

We then tested whether the other four datasets had similar performance. First, we observed that LDA models achieved high classification performance on the held-out test set for all datasets (see Table S6 for accuracy, precision, recall and F_1_ score) and were not overfit as evidenced by similar performance on the training and test sets. Next, we observed that LDA projections were highly correlated with continuous measurements, as evidenced by Spearman’s ρ between 0.5 and 0.85 (**Figure 3B**, additionally see Figures S1-5 for hyperparameter effect on performance). To get an estimate of model sensitivity to dataset splitting, we performed 5-fold cross validation (see Methods) on each training dataset (Figure S6). Strikingly, for each of the datasets, we observed no significant difference in the correlation (significance was measured as a t-test on the unbounded Z transform of the Spearman ρ).^40^ Encouraged by the success of correlation, we also sought to explain the magnitude of correlation, which was consistently high but had two outliers. **Adams 2016** dataset had a significantly lower predictive value of ρ∼0.5. We suspect this decrease in performance has two sources: noise in the dataset due to an abundance of unresolvable low affinity variants (see Figure S8 for correlation plots for each dataset), and the lack of discrimination between binding affinity and expression level in the experimental sorting design, which can attribute higher affinity to sequences with higher display and vice versa.^17^ The **Makowski 2022** specificity dataset also had lower than average performance; we hypothesize this model suffered due to the difficult nature of measuring antibody off-target binding.^27, 41, 42^

To test whether linear models were limiting the predictive capabilities of continuous properties, we also tested fully connected, feed forward neural networks, which have been shown to similarly identify continuous values from binary data.^27^ While non-linear models may capture higher order epistatic behavior, these models generally performed as strongly as LDA models (Figure S7). Over this wide range of protein engineering objectives, this approach consistently predicts continuous properties and has comparable accuracy to models trained on state-of-the-art sequencing and sorting data.

### Prediction of stapled peptide affinity and specificity from binary labels

To apply this method prospectively to a new dataset following the promising retrospective analysis, we chose B cell lymphoma 2 (Bcl-2) stapled peptide antagonists as our design case. In addition to requiring non-natural amino acids, making it incompatible with modeling approaches based on naturally evolved proteins, these peptides are well suited for this approach because we can evaluate not just a single property but the tradeoff between affinity and specificity. We generated a dataset of B cell lymphoma 2 (Bcl-2) stapled peptide variants that were sorted over several rounds **(Figure 4A)** using the bacterial cell surface display.^31, 32^ This library was designed based on naturally occurring peptide sequences, SPOT arrays of BIM mutants, and previously designed high-affinity or specificity BH3 variants (Case 2023, manuscript in progress) (see Table S2-4)^24, 43, 44^ Because bacterial surface display libraries are highly limited by size compared to the theoretical diversity of BH3 peptides (∼10^30^), mutations were prioritized that were predicted to govern specificity between Mcl-1 and Bcl-x_L_. The final library of ∼10^9^ was transformed into bacteria (**Figure 4B**) and sorted against either Mcl-1 or Bcl-x_L_ with a combination of three magnetic and four fluorescent activated cell sorting (MACS/FACS) (**Figure 4C**). The magnetic cell sorting was performed until the library was sufficiently reduced in diversity for analysis with FACS, which offers more precise control over property selection. We deep sequenced these pools to isolate highly active peptides (**Figure 4D**), which enabled an understanding of sequence trends that governed high affinity and specificity (see Figure S9 for sequence trends) and provided a source of data to train and evaluate the capabilities of LDA models to predict peptide function (**Figure 4E-F**). We observed high correlation between for both Mcl-1 and Bcl-x_L_ LDA models (Spearman’s ρ of 0.893 for Mcl-1 and 0.708 for Bcl-x_L_).

**Figure 4:**
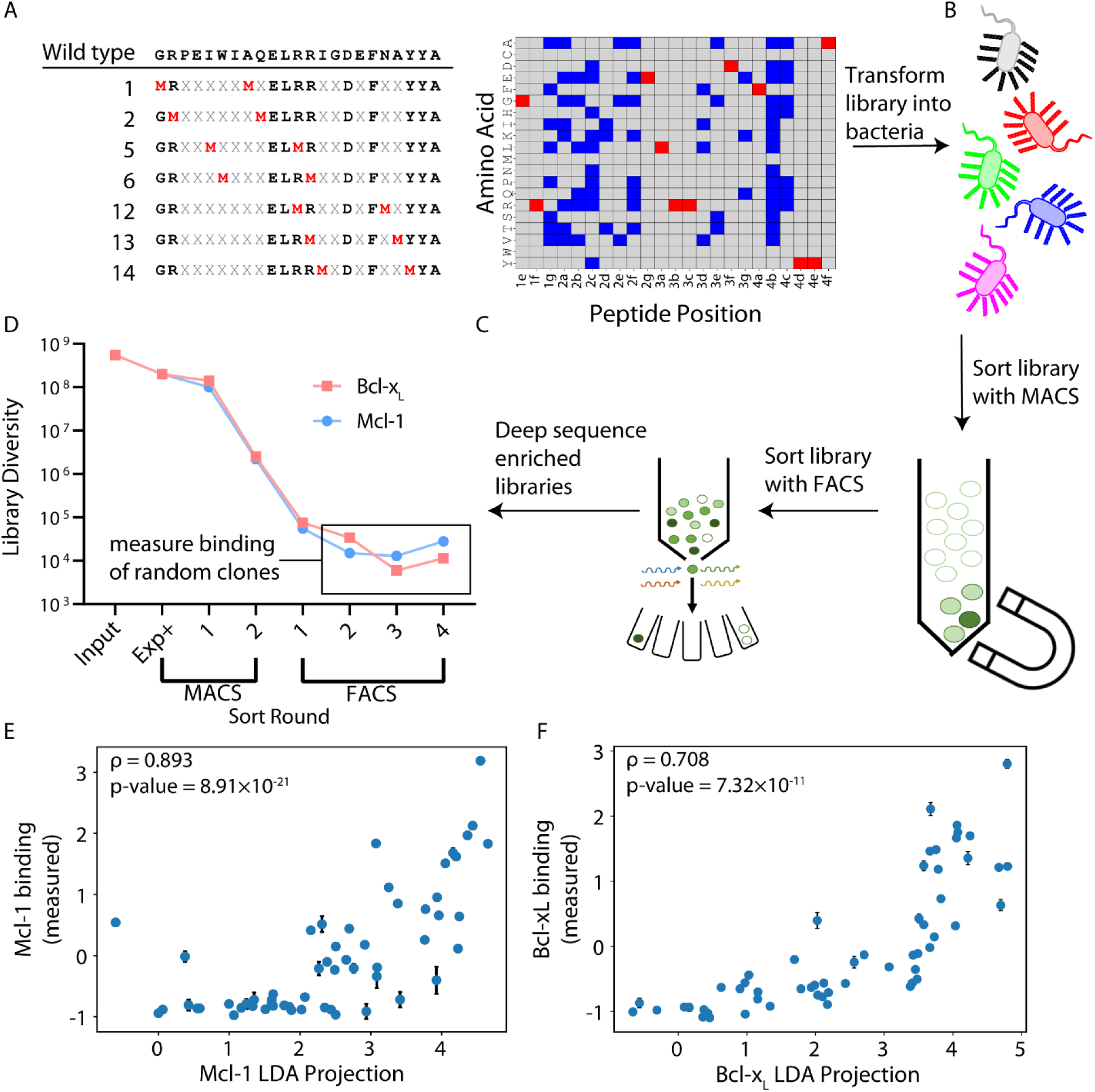
Prospective analysis of B cell lymphoma 2 (Bcl-2) pro-apoptotic stapled peptides via bacterial surface display, deep sequencing, and machine learning. A combinatorial mutagenesis library of stapled BIM variants was designed including staple locations (left) and sequence (red positions fixed, blue positions variable, right) **(A)**, transformed into bacteria **(B)**, sorted using a combination of magnetic activated cell sorting (MACS) **(C)** and fluorescent activated cell sorting (FACS) towards Bcl-xL and Mcl-1 (two members of the Bcl-2 family) in parallel. The library was next generation sequenced (NGS) to calculate frequencies of each unique sequence along the sorting progression **(D)**. Finally, a LDA model was trained on the binary labels from NGS and used to predict the continuous binding of 57 peptide variants, which were selected randomly from FACS 2-4 for both Mcl-1 **(E)** and Bcl-xL **(F)**.

To generate training data, we aggregated all four rounds of FACS and the expression positive MACS sorts, hypothesizing that would provide additional confidence for ‘hits’ and expressing but non-binding sequences. The ratio of these counts was computed as described above and used to generate labels for LDA training and testing (see Figure S9 for logoplots of negative and positive sequences). First, we observed that LDA models had high classification performance and were not overfit (see Table S7 for performance statistics and Figure S10 for hyperparameter effect). We then tested the performance of LDA to predict continuous properties by randomly sampling 57 sequences among the FACS sorts, measuring their continuous binding via flow cytometry, and measuring the correlation between predicted LDA binding and the sequences’ actual binding **(Figure 4E-F)** (see Figure S11 for sequences and data). We observed a strong correlative power between LDA projections and continuous measurements of peptide affinity: spearman ρ of ∼0.7 and ∼0.8 for Mcl-1 and Bcl-x_L_ respectively (p < 0.00001). Finally, we sorted the final round of sorted cells via SORTCERY for a comparison with high-throughput, semi-quantitative measurements of binding affinity. Surprisingly, the binary sorting data coupled with an LDA model trained with NGS data had better performance than selecting clones from the final 2 rounds of sorting for Mcl-1 specificity (Figure S12), suggesting that the information contained from simple sorting experiments provides a powerful method to predict continuous protein properties.

### Optimization of stapled peptides using machine learning and integer linear programming

While directed evolution campaigns may yield the desired properties after sorting, sequencing, and modeling, it is also possible that further optimization is necessary. In such cases, protein engineers rely on a combination of manual and automated approaches to further optimize lead candidates.^24, 34–36^ We sought to explore how our modeling workflow could not only score entire sequences, but how the contributions of individual amino acids contributed, potentially enabling the generation of new, unsampled sequences. Because linear models have associated weights for each amino acid and sequence position, the same scoring tools to find the best measured clones can also be used to score sequences that have never been evaluated experimentally. We therefore applied an optimization approach that can optimize discrete inputs for continuous properties and explore unseen sequence space: integer linear programming (ILP) **(Figure 5)**, which has previously been applied to design specific linear peptides towards the Bcl- 2 proteins.^24^ To establish the baseline of specificity from sorting, we further characterized variants from the final round of sorting that were predicted to be specific for Bcl-x_L_ and Mcl-1. Interestingly, most peptides from the Bcl-x_L_ library were highly specific (Figure S13), while fewer from Mcl-1 performed favorably (∼80% had significant off-target binding, **Figure 5A**). We hypothesized we could recover specific Mcl-1 clones by optimizing sequences from sorting and sequencing data that otherwise yielded mixed results. We solved the ILP model three times, once for Bcl-x_L_ specific peptides, once for Mcl-1 specific peptides, and once more for bispecific peptides (see Methods for more details). Out of many sequences predicted to have high activity for Mcl-1 (Figure S14), we randomly selected XXX sequences for low-throughput flow cytometry analysis (Table S8). Strikingly, we observed that the optimized Mcl-1 sequences displayed similar or improved specificity compared to the highest activity clones assayed experimentally.

**Figure 5:**
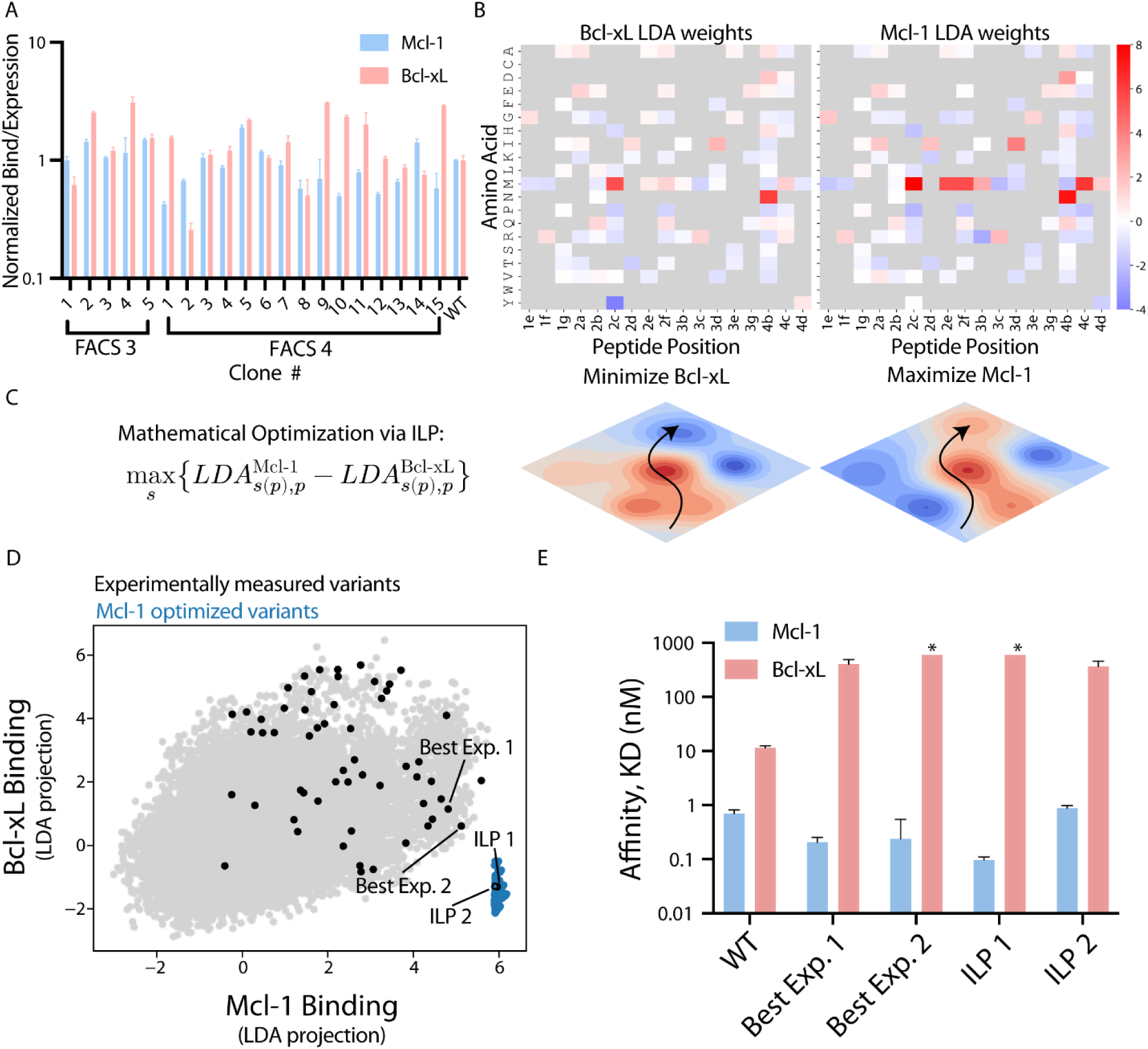
Extrapolation of interpretable ML model weights to generate novel, highly specific Mcl-1 inhibitors. Of 20 sequences randomly selected from the final two rounds of sorting towards Mcl-1, many did not display high levels of specificity towards Mcl-1 when measured in low throughput binding assays **(A)**. We hypothesized the weights from linear discriminant analysis (LDA) machine learning could be used to design peptides with high affinity to Mcl-1 **(B)** or Bcl-xL **(C)**. To optimize the sequences, we applied integer linear programming (ILP) **(C)** to maximize the likelihood a peptide binds Mcl-1 while minimizing its binding to Bcl-xL **(D)**. ILP identified numerous sequences that were predicted to be highly specific **(E)** that were not among the 10^5^ sequences assayed experimentally. Two variants were randomly chosen among this set and were found to be as specific as the best clones identified from sorting **(F)**.

Sequences initially identified by minimizing Mcl-1 binding while maximizing Bcl-x_L_ binding resulted in peptides that did not bind either Mcl-1 or Bcl-x_L_ (Figure S15 and Table S8); it has been previously shown that subtle differences in ILP set up can affect the efficiency of outcome.^24^ We suspect this failure was due to the model being overly sensitive to mutation at Asp at position 4b, which was the only mutation consistently sampled that had a high score for both Bcl-x_L_ and Mcl-1 but was slightly higher for Mcl-1. To address this issue, we maximized Bcl-x_L_ binding then chose the sequences which had the lowest Mcl-1 scores, which preserved Bcl-x_L_ binding and resulted in highly specific peptides (Table S9 and Figure S16). While our sorting campaign was originally designed to identify highly specific peptides, we also pursued bispecific peptides, which serve as proof that the model can interpolate in sequence-function space but could also serve as therapeutics in diseases driven by both Bcl-2 proteins. Sequences were identified by maximizing both Mcl-1 and Bcl-x_L_ binding, yielding peptides with relatively high affinity for both targets that had significant sequence difference from wild type (BIM) (Figure S17).

To show generalizability of ILP to generate functional protein variants, we additionally set up the optimization problem using the **Makowski dataset** (Figure S18). We defined the objective of this optimization as the minimization of off-target binding, subject to the maintenance of affinity. We solved the model and compared the highest functional sequences according to our predictions to those described in the original manuscript. We found that the predicted sequences were extremely close to those identified as co-optimal by Makowski and co-authors.

## Discussion

In this work, we developed a method to utilize NGS data from simple binary sorting results with machine learning to infer continuous protein properties. These results can also be utilized to extend the sequence space beyond sequences directly observed in the library **(Figure 1)**. The workflow consists of two important parts: the label assignment process from deep sequencing data, and the use of linear machine learning models to predict continuous protein properties from binary data **(Figure 2)**. Currently, there is a lack of consensus on how to best analyze directed evolution data for lead molecule selection and protein optimization. This lack of consensus likely arises from variations in how experiments are set up, which depends on surface display platform, sequencing instrumentation, FACS instrumentation, the design of sort gates, sequencing depth, among other factors. This technique provides a practical but powerful method compared to typical enrichment ratio analysis through a simple binary classification from any sorting experiment. By defining a ratio of frequencies based on any two gates (positive/negative sort, input/output sort, etc.) and binarizing the ratios into ‘1’ and ‘0’, any directed evolution experiment can be transformed into a dataset for downstream analysis. The transformation to binary labels is important because the next component of the workflow is the use of linear machine learning models that can be used to predict continuous properties from directed evolution data (linear discriminant analysis, LDA).^27^ The noise in enrichment ratios is likely mitigated by binarization, and the information contained from labels and sorted protein sequences facilitates the continuous transformation yielded by machine learning models.

To test our method, we curated data from five large protein engineering campaigns: the fluorescent landscape of avGFP,^28^ the directed evolution of a fluorescein-binding scFv,^21^ the RBD affinity landscape towards SARS-COV-2 Spike protein,^25, 26^ high-affinity and high-specificity Fabs,^27^ and the design of highly specific peptides against B cell lymphoma 2 (Bcl-2) proteins **(Figure 3)**.^24^ Proteins in these data vary in complexity from short alpha helical peptides to large globular proteins and in objective from protein fluorescence to multi-objective affinity and specificity optimization. Furthermore, each of these datasets varied in both sorting strategy and complexity: **Makowski et al.** sorted for the top ∼5% of antibody variants while **Adams et al.** quantified the binding of an entire family of fluorescein binders. While many of the projects relied on complex sorting techniques to obtain quantitative protein labels, we simulated simple binary sorting experiments by limiting the sequencing data (see Methods). We then evaluated the predictive power of LDA models trained on these simple sorting experiments and observed both impressive classification performance and strong prediction of continuous properties from LDA binary projections. Interestingly, models trained on binary data were highly correlated with continuous data (Spearman correlation coefficients ranged from 0.5-0.9). Furthermore, when we compared the predictive power of LDA models trained on binary data to regression models trained on continuous data, we observed no increase in rank order performance, suggesting that models trained on simple sorting experiments yield comparable information to models trained on data from experiments that generate hundreds to thousands of continuous measurements.^21, 23, 24^

Next, we sought to explore how this workflow could be used for prospective analysis in addition to retrospective analysis **(Figure 4)**. We hypothesized that because the workflow is agnostic to protein type and display platform, any directed evolution campaign with sufficient sorting and sequencing data is a suitable environment for testing. As such, we chose to analyze libraries of stapled peptides, an important class of protein formed by a covalent crosslinking of two amino acids.^45^ Stapled peptides are being explored as therapeutics for previous ‘undruggable’ disease related proteins, owing to their location inside the cell and untargetable by small molecule drugs.^46^ Stabilized Peptide Engineering by *E. coli* Display (SPEED) has previously been demonstrated to accelerate the development of stapled peptides by displaying them on the surface of bacteria, where libraries of peptides varying in sequence and staple location simultaneously can be optimized for protein-peptide interactions.^31, 32^ One additional challenge in the optimization of stapled peptides is their reliance on non-natural amino acids, which generally results in the incompatibility of models trained on naturally occurring sequences.^47–50^ We built on previous work by generating a library of randomized stapled peptides towards two B cell lymphoma 2 proteins (Bcl-2), an important class of apoptosis regulatory proteins that is responsible for cancer cells immortality.^51^ We sorted this library against two important members: Mcl-1 and Bcl-x_L_,^52^ each of which drives immortality in different diseases.^53^ Selective targeting among Bcl-2 proteins is an outstanding goal in drug targeting but is difficult due to the highly homologous nature of these proteins. After several rounds of cell sorting and subsequent deep sequencing, we trained LDA models on a subset of the binary sequencing data, evaluated the model on both the hold-out test set, and generally observed high classification performance. We then measured the binding of 57 sequences from various rounds of sorting with low throughput flow cytometry experiments and observed that many of the clones did not demonstrate favorable affinity or specificity properties when sampling from these enriched libraries. However, we did observe a high degree of correlative power between LDA projections and continuous peptide binding. Finally, these models were able to identify molecules within the set of experimentally observed sequences that were highly specific but may not have been selected for lead compounds due to their rarity.^11^ Several sequences along the Pareto frontier, or the boundary of co-optimality where an increase in one property leads to a decrease in the other, were translated into bacteria and assayed via flow cytometry. We also characterized several clones that were bispecific, which could have applications in specific diseases, but also serves as a test case if the model can interpolate function where it wasn’t directly engineered via cell sorting. The specificities of these peptides agreed with model predictions, indicating the model was able to identify functional and rare peptides from across the specificity landscape.

Finally, we sought to use the interpretive nature of the linear machine learning models to explore unseen sequence space and generate highly diverse and novel sequences **(Figure 5)**. To accomplish this task, we used integer linear programming and the coefficients from machine learning to mathematically optimize peptide sequences beyond the properties that were experimentally observed (from deep sequencing or flow cytometry).^24^ We hypothesized that such an approach could recover functional peptides with consistency where sorting did not; while the final round of Bcl-x_L_ sorting yielded consistently high affinity and specificity variants (Figure S13), the Mcl-1 sort had a small fraction of sequence variants with desired properties (Figure 5A). We thus prioritized the design of Mcl-1 binders and identified a new peptide sequence that improved peptide properties beyond the experimentally measured Pareto front. Importantly, this variant demonstrated specificity at least as potent as the most specific clone identified from experimental work.

To test whether our sequence optimization workflow generalizes beyond small alpha helices, we also applied ILP to the **Makowski dataset** (Figure S18). While antibodies have been the subject of optimization using highly sophisticated models,^54–57^ we hypothesized that the high performance from linear ML models would make it amenable to ILP optimization. Like Bcl-2 inhibitors, antibodies need to demonstrate properties beyond high affinity to be considered therapeutic, and ILP is uniquely suited to tackle co-optimization.^58^ We observed that the set of sequences predicted to be co-optimal by ILP are similar to their most optimal clones identified experimentally. Furthermore, their lead antibody identified as co-optimal (EM1) was among the set of antibodies predicted by ILP. Makowski and co-authors designed a comparatively small library (∼10^6^) for their experimentally measured sequences (∼10^4^), resulting in a more confident sampling of mutated amino acids experimentally. In contrast, the library of stapled peptides we designed had a much larger ratio of design space (∼10^9^) to experimentally measured sequences (∼10^5^), making this library suitable for extrapolation beyond experimentally measured space using machine learning. For protein variant libraries where mutations are sufficiently independent (minimal higher order epistatic interactions), a strategic subsampling of design space can be advantageous for subsequent protein optimization with linear models^59, 60^ and help to de-risk sorting campaigns, as exploration through the full design space can improve function beyond those originally assayed.

The use of ML with NGS data from binary sorting campaigns has many advantages, but the approach also has a few limitations. It is important to note that LDA projections are correlated with, but not predictive of, continuous measurements. Therefore, LDA-informed properties may not match 1:1 with continuous properties. However, because many protein engineering campaigns do not seek to quantify the exact magnitude of fitness, but rather seek to maximize or minimize a property or trade-off between properties, this correlation can still provide direct insight into protein fitness and accelerate optimization efforts. We also found that ILP optimization was sensitive to model weights as evidenced by the initial failure of generating highly specific Bcl-x_L_ peptides. Two approaches to address this are incorporating uncertainty into model predictions that could yield more confident extrapolation into unseen sequence space,^34^ or selecting a range of sequences from multiple modes of optimization simultaneously.^24^ Despite identifying peptides with high specificity towards Mcl-1 and Bcl-x_L_, more work is needed to yield effective peptide therapeutics: it is equally important to show these peptides do not bind the other 3 Bcl-2 members.^53^ Lacking knowledge of the sequence space of high affinity binders, we were unable to explore this aspect of peptide design; future work includes designing stapled peptides against the entire Bcl-2 family, which play additional roles in off-target toxicity and are responsible for immortality in other cancers. Because this approach is amenable to higher dimension multi-objective optimization, we expect that optimizing specificity for five proteins with this approach is possible.

Despite these limitations, the ability to score sequences beyond those observed experimentally is important because drug-like properties not easily assayable by high-throughput techniques (immunogenicity, stability, cell permeability, etc.) are often highly dependent on sequence and may need further optimization.^58, 61–63^ For example, minimization of positive charge in CDR regions of antibodies has been shown to minimize off-target binding,^42^ while selective placement of hydrophobicity and positive charge has been shown to improve cell penetration for stapled peptides.^62, 64^ This combined machine learning and optimization approach provides a powerful method to identify highly functional protein variants if experimentally measured clones did not meet fitness criteria or further sequence optimization is necessary.

In summary, the data processing and modeling workflow designed in this work is a versatile tool towards the improved analysis and identification of protein variants across many domains of protein engineering by utilizing machine learning and NGS data to predict continuous properties from binary sorting data.

## Notes

### Competing Interest Statement

The authors have declared no competing interest.

