## Supporting Figures 1-18 and Tables 1-9 for "Machine Learning to Predict Continuous Protein Properties from Simple Binary Sorting and Deep Sequencing Data"

^1^Chemical Engineering and ^2^Biomedical Engineering

University of Michigan, Ann Arbor, MI 48109, USA

Keywords: protein engineering, directed evolution, supervised machine learning

Running title:

**Table S1:** Parameters of each dataset used in this study

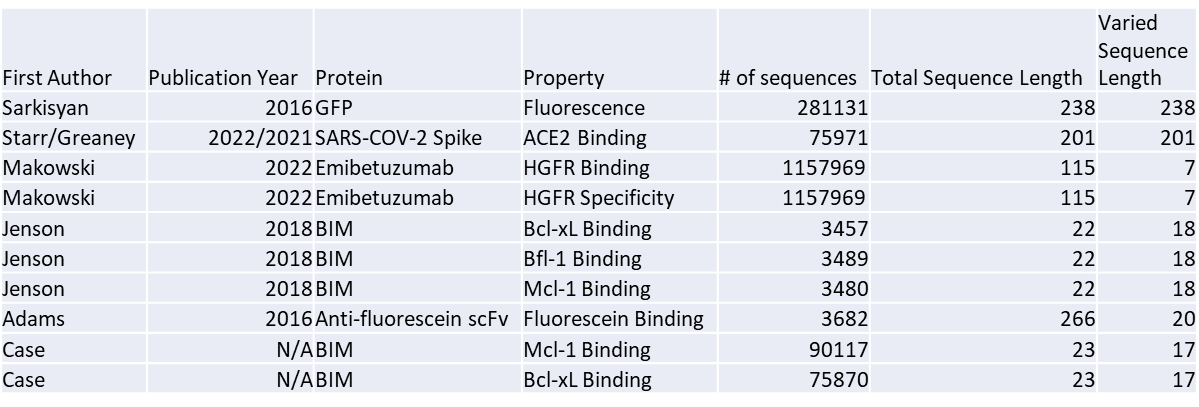

**Table S2:** Degenerate codon design for pro-apoptotic anti-Bcl-2 bacterial surface display library

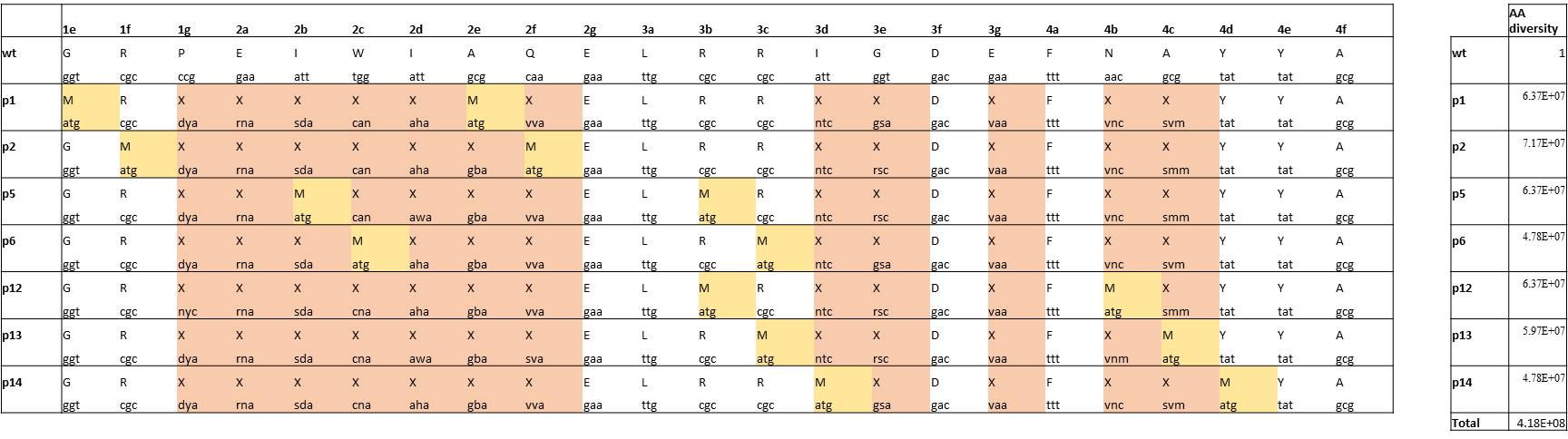

**Table S3:** Sampled amino acids for pro-apoptotic anti-Bcl-2 bacterial surface display library

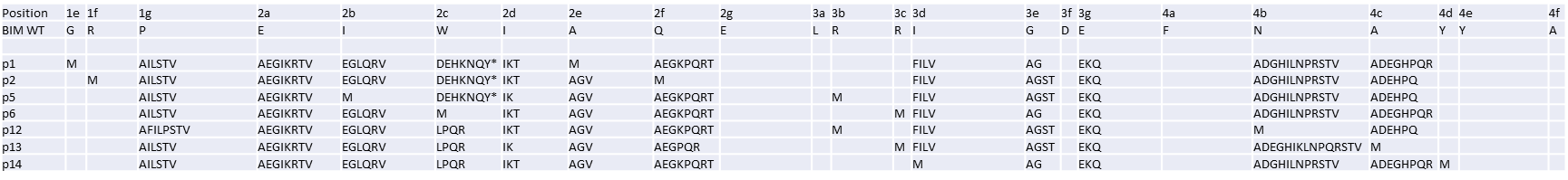

**Table S4:** Library design primers for pro-apoptotic anti-Bcl-2 bacterial surface display library

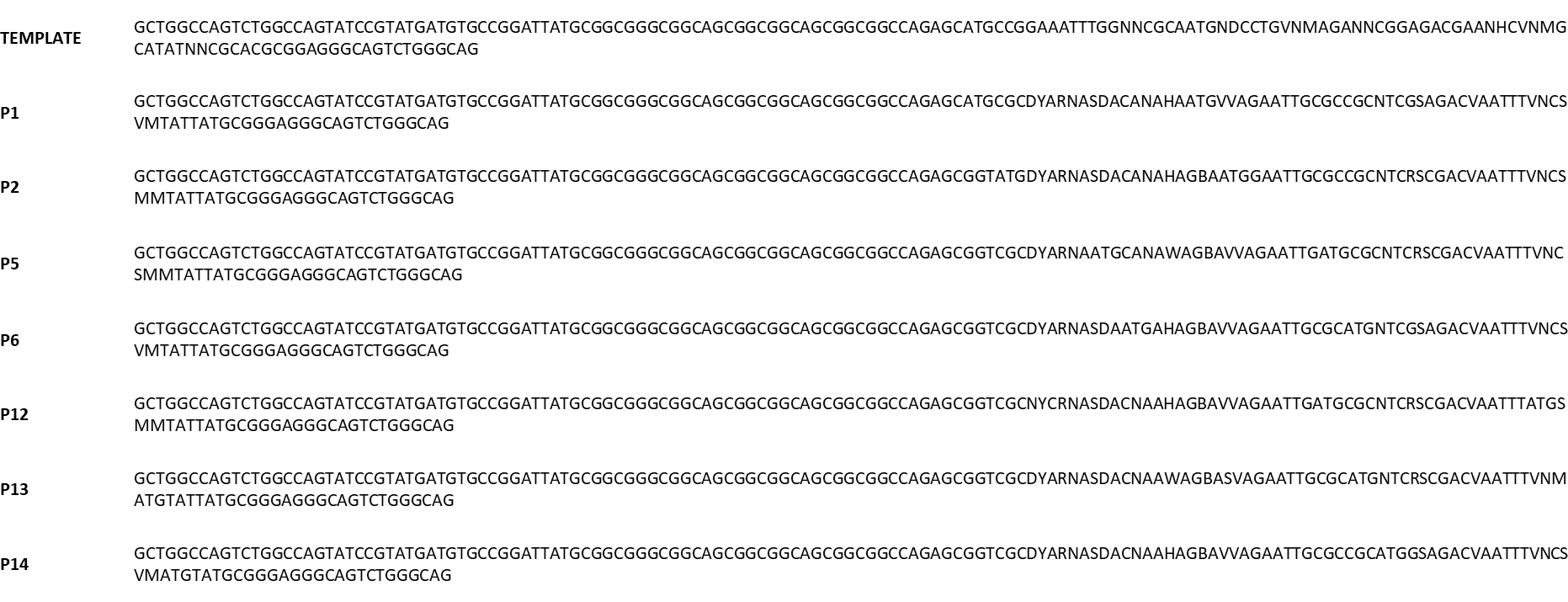

**Table S5:** Next generation sequencing primers for bacterial cell surface display

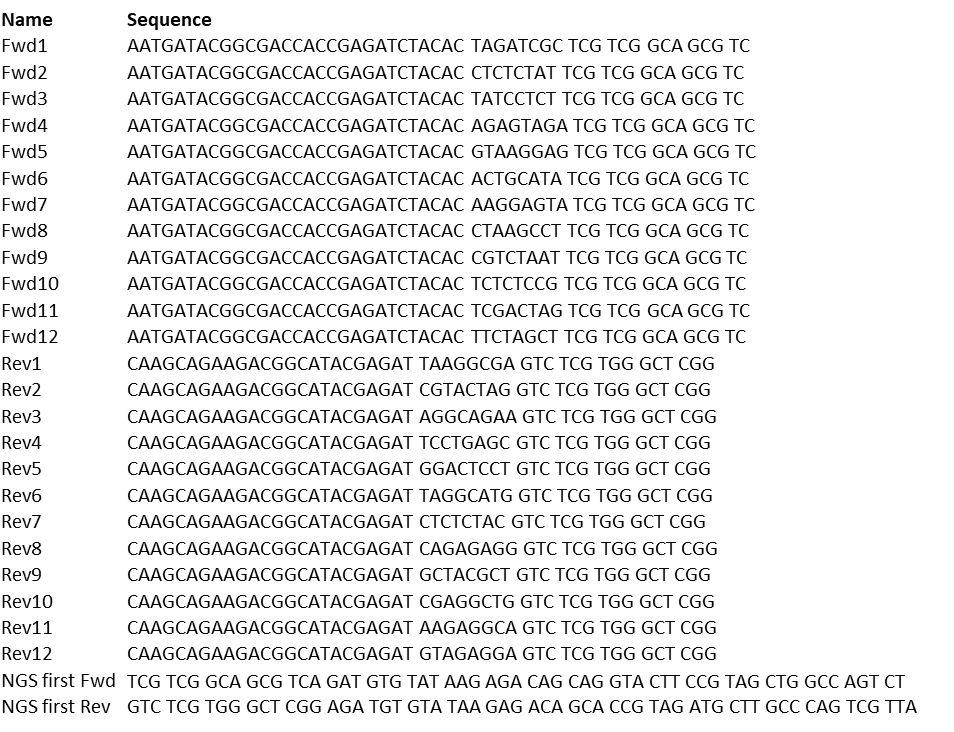

**Table S6:** Linear discriminant analysis classification performance for previously reported datasets

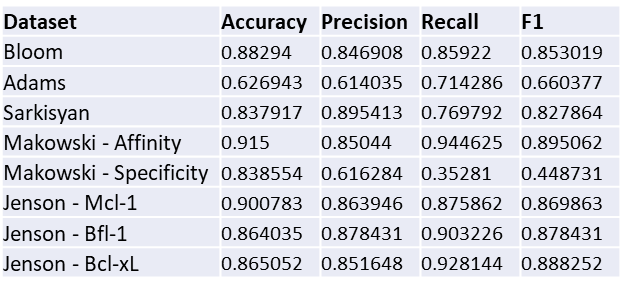

**Table S7:** Linear discriminant analysis classification performance for stapled peptide library

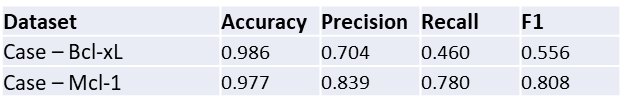

**Table S8:** Integer linear programming designed sequence and primers

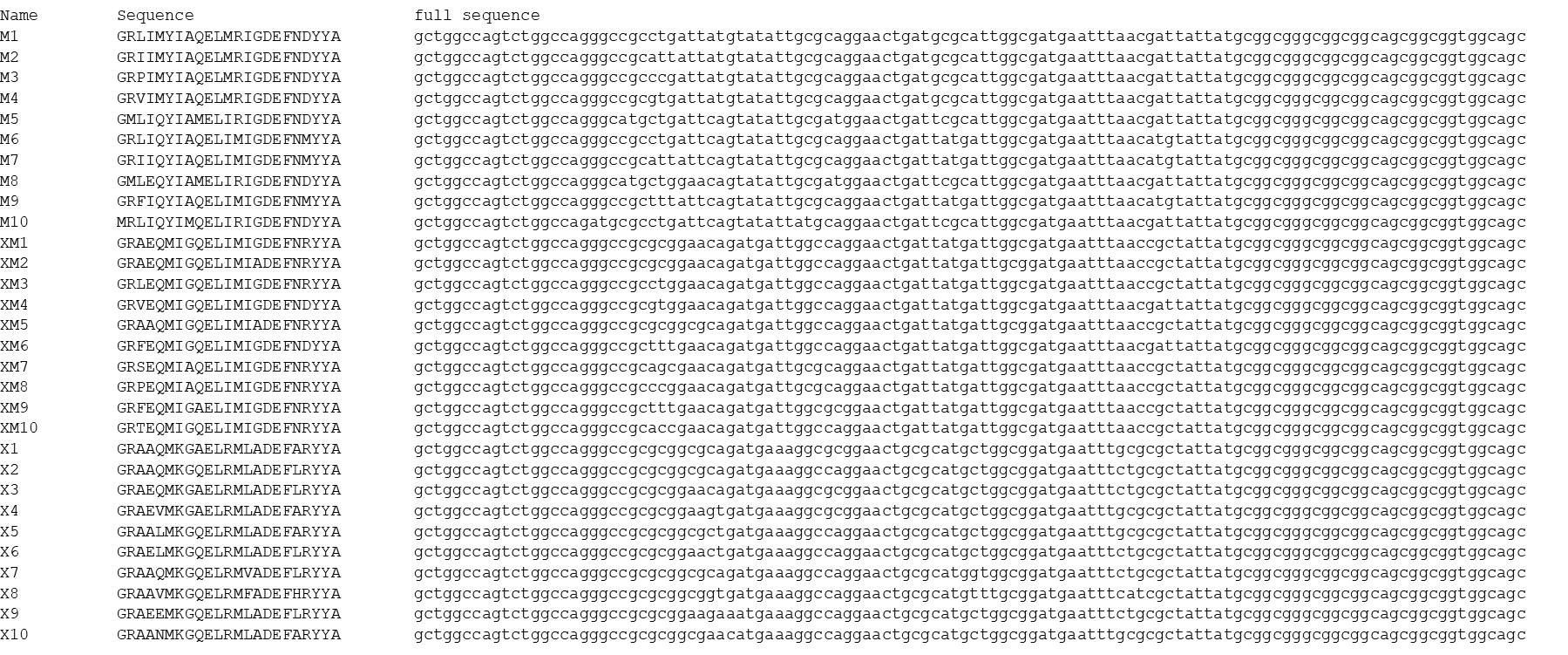

**Table S9:** Integer linear programming designed sequences for Bcl-xL design 2

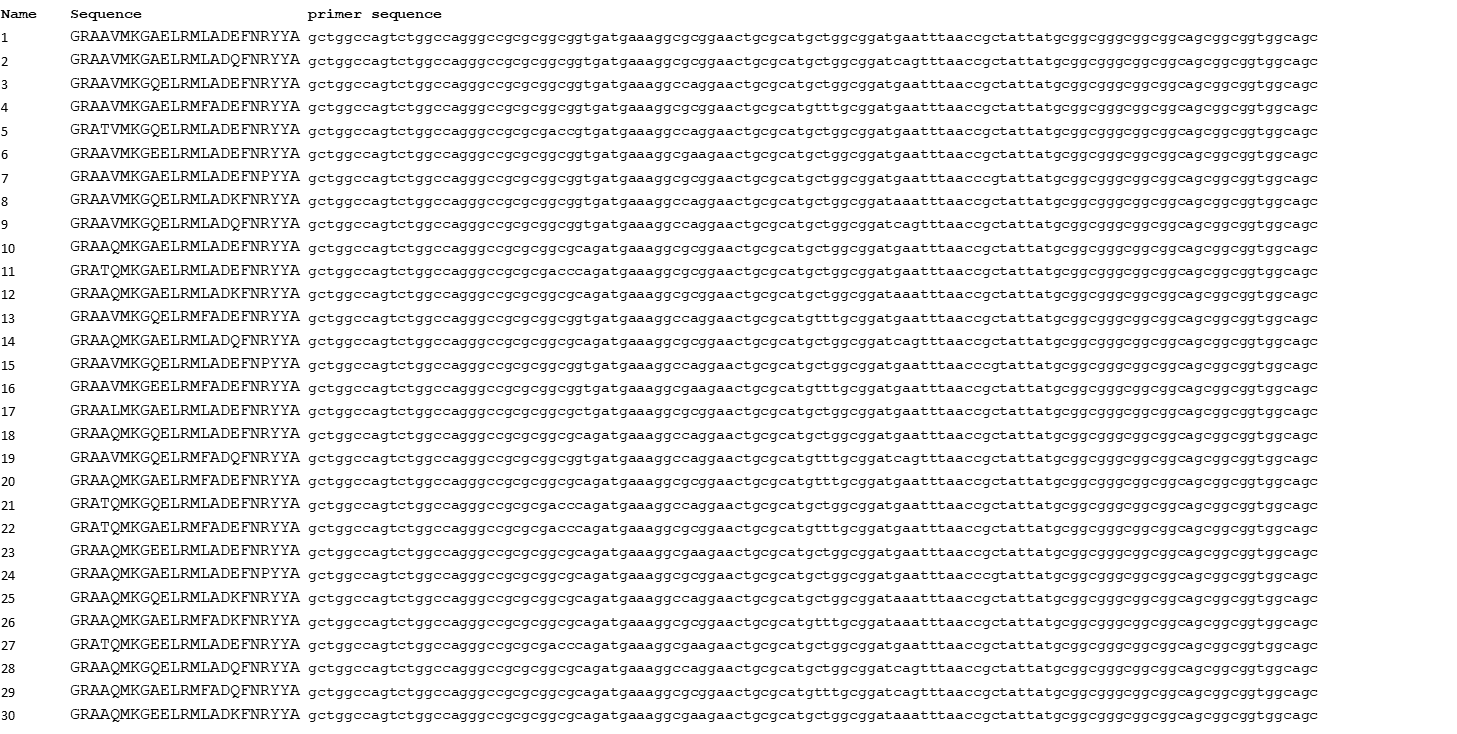

**Figure S1: Dataset hyperparameters for Makowski et al. (2022) Nature Communications.**

**Makowski 2022 - Affinity**

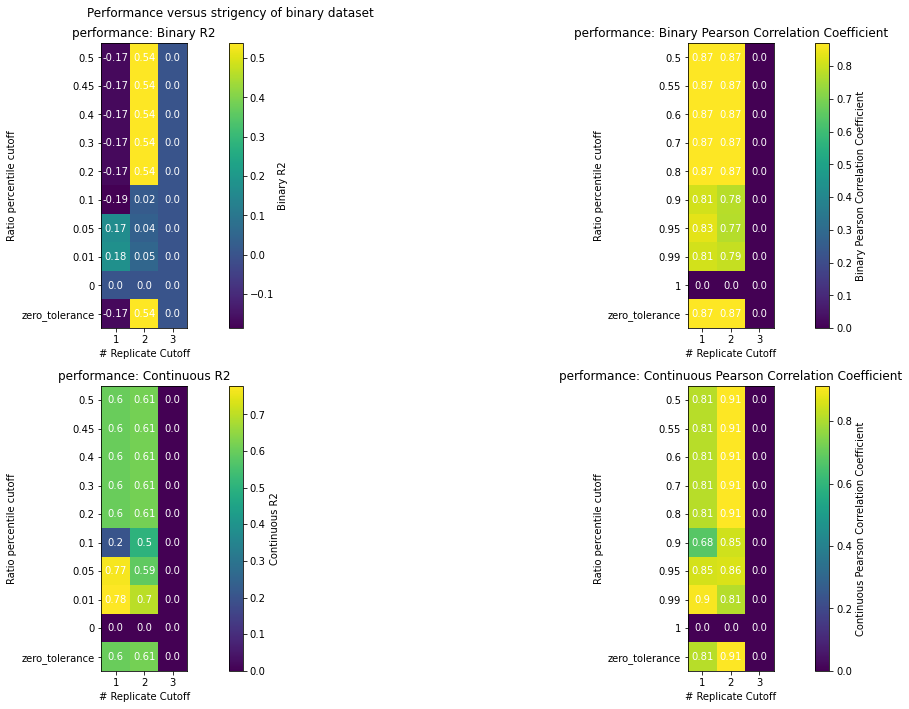

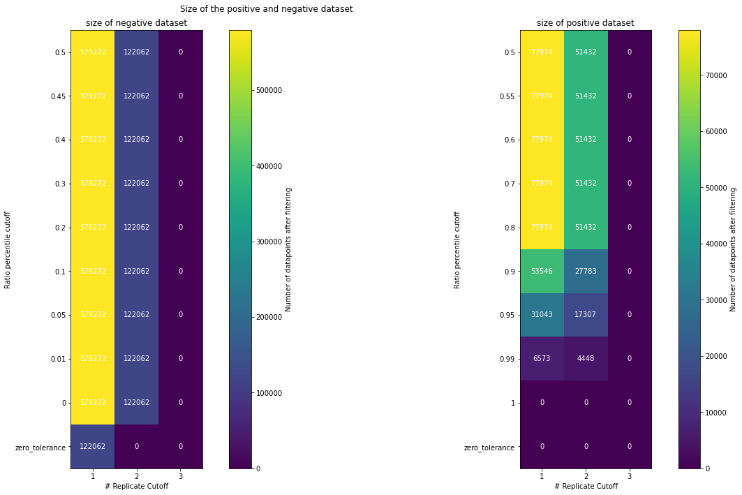

**Makowski 2022 - Specificity**

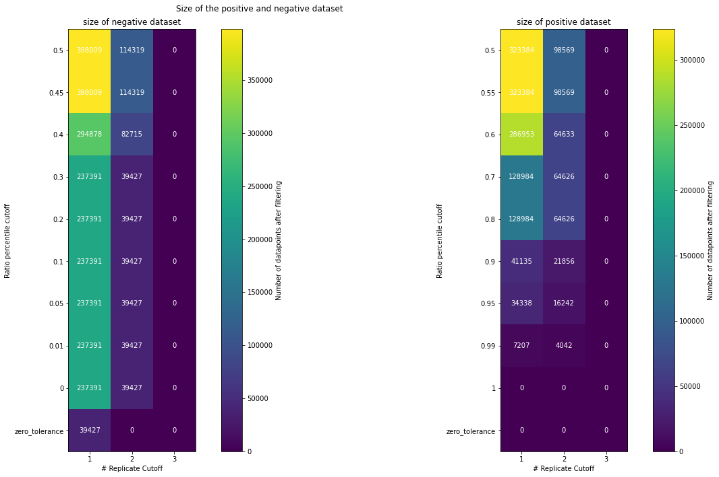

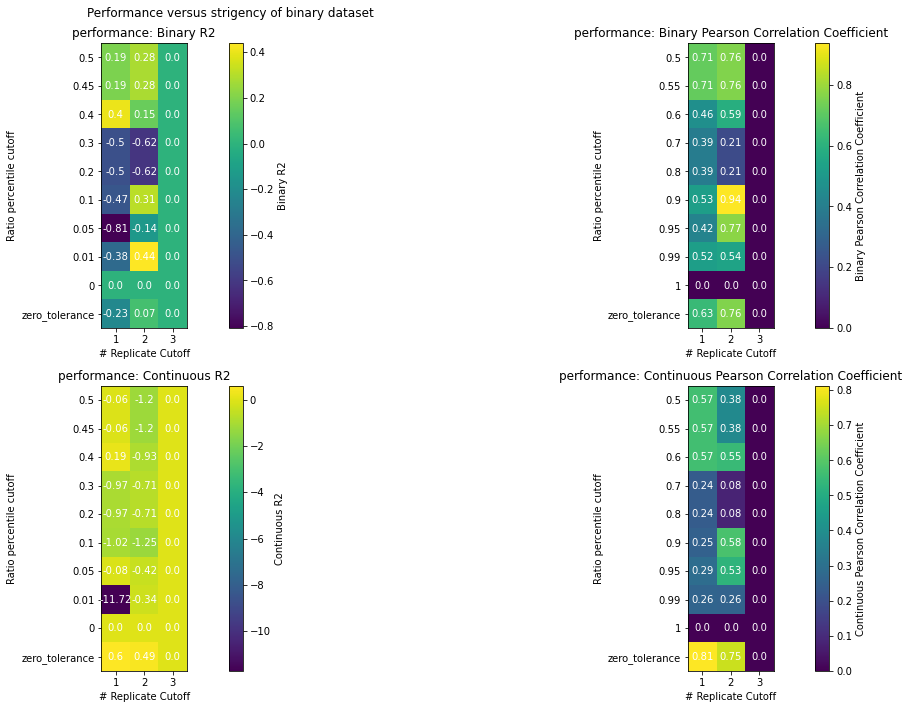

**Starr 2022**

**Figure S2: Dataset hyperparameters for Starr et al. (2022) Science.**

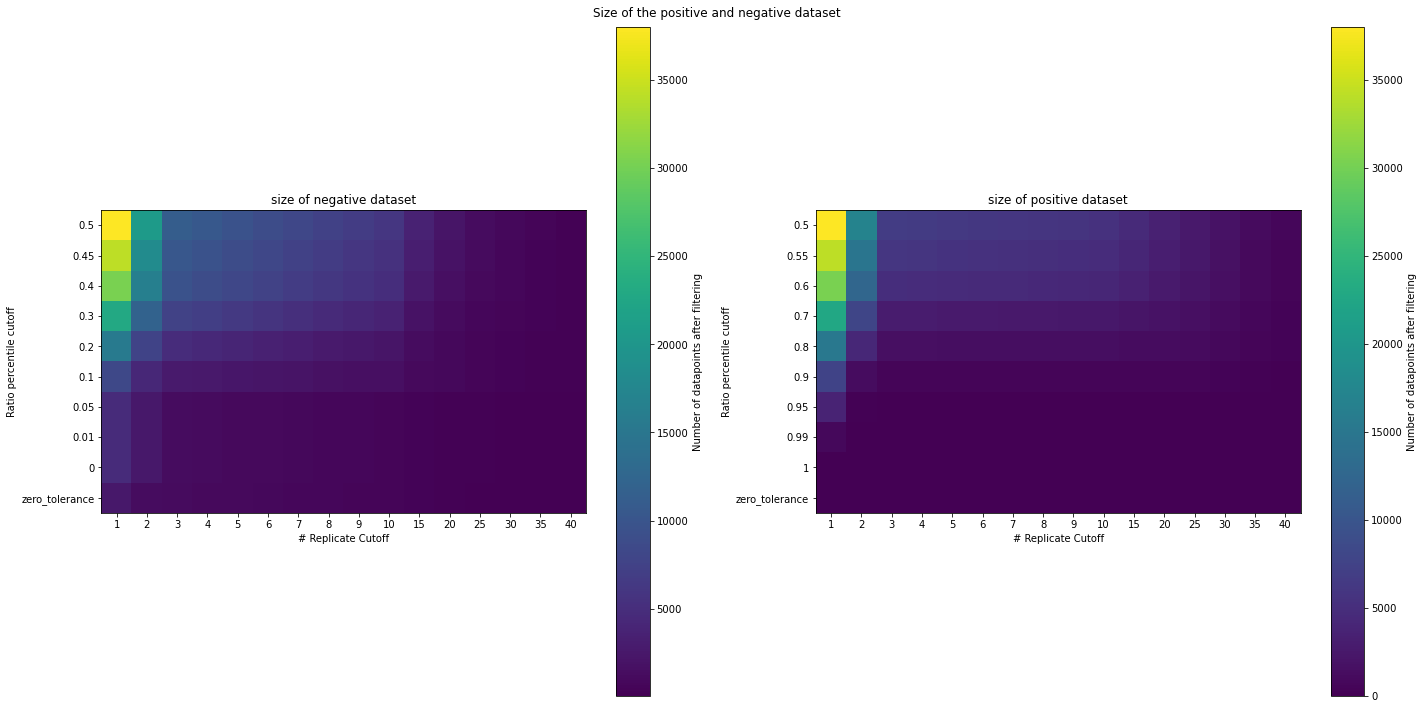

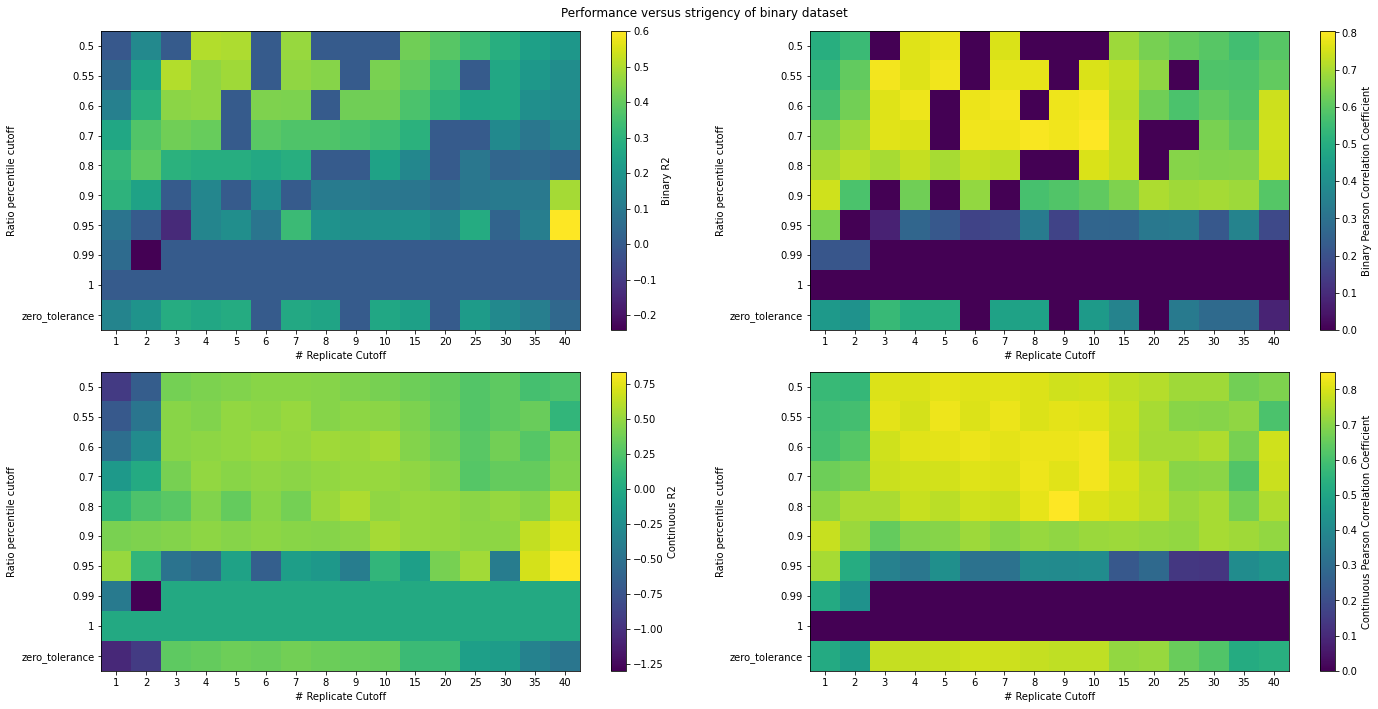

**Figure S3: Dataset hyperparameters for Sarkisyan et al. (2016) Nature.**

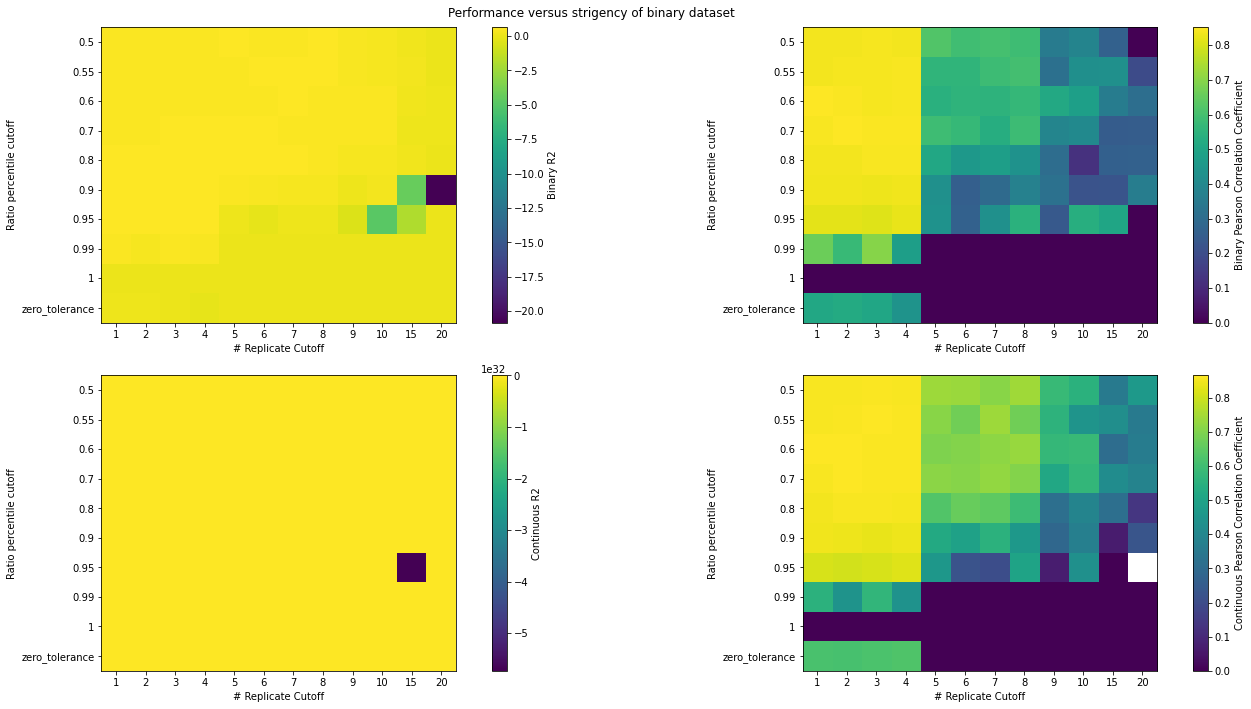

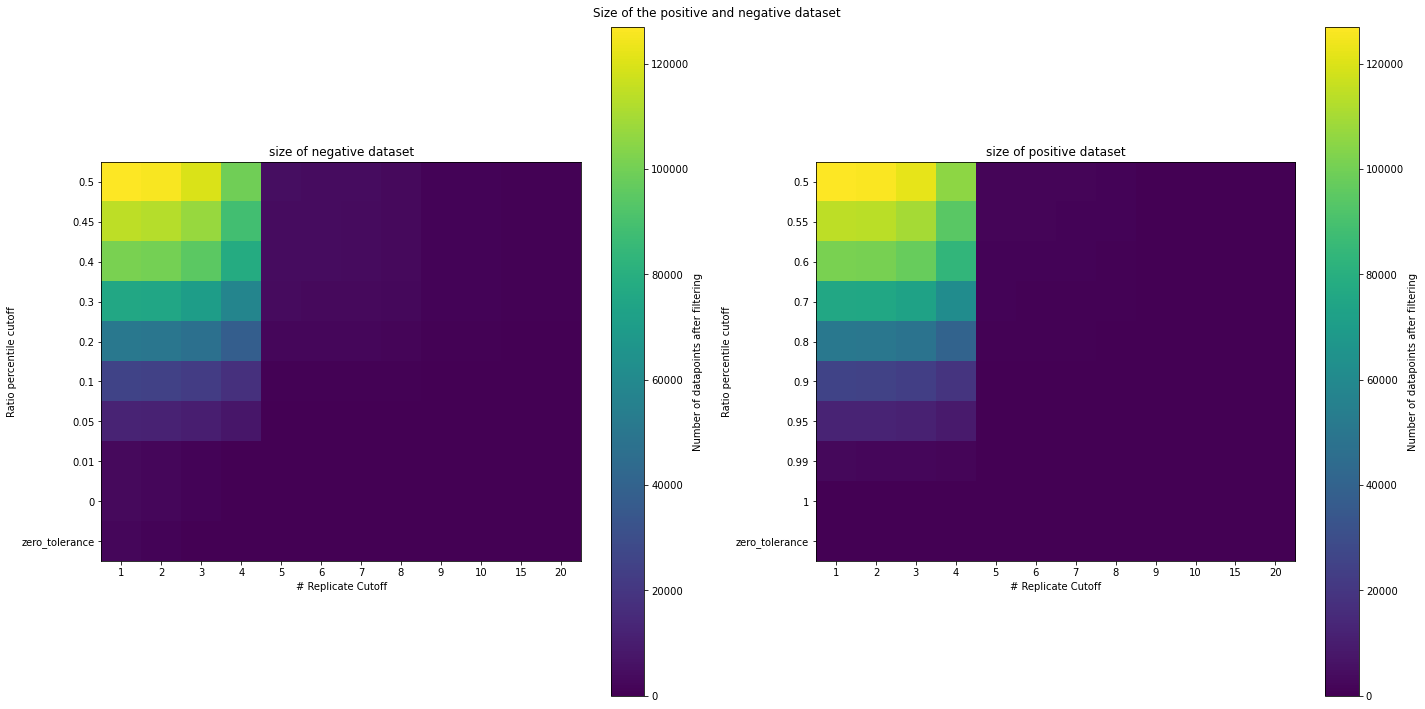

**Sarkisyan 2016**

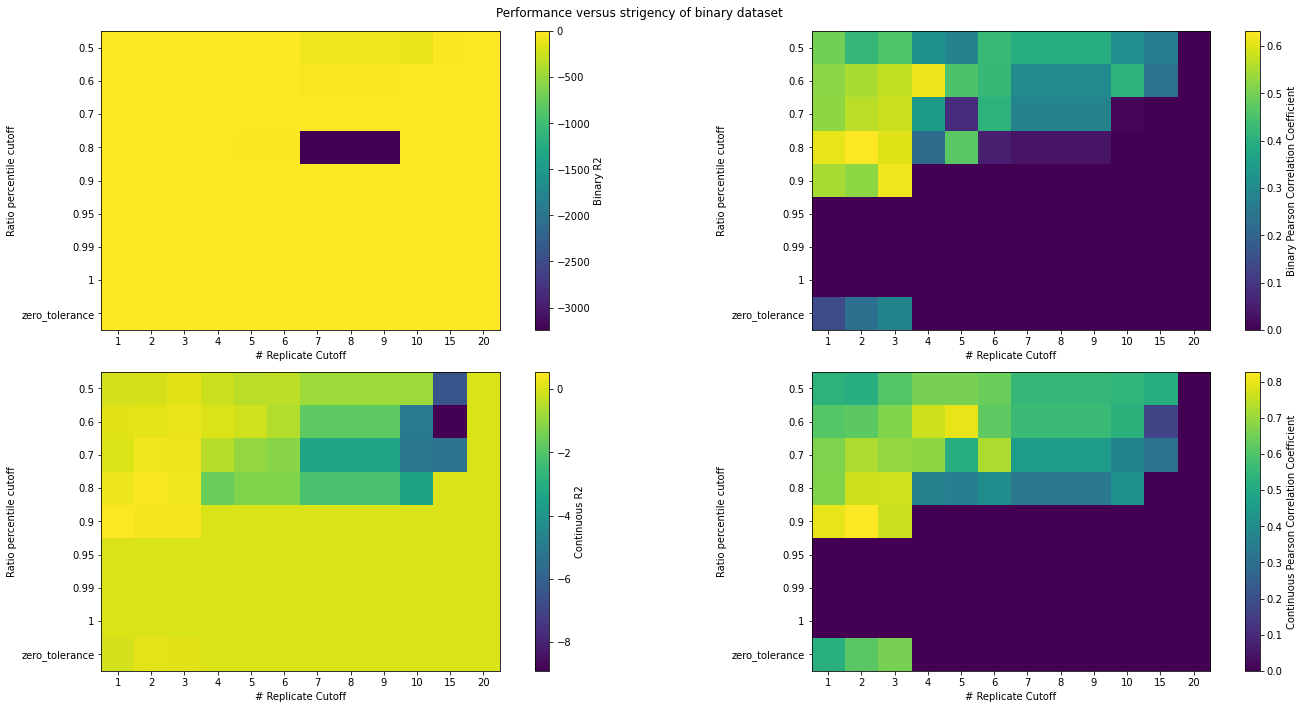

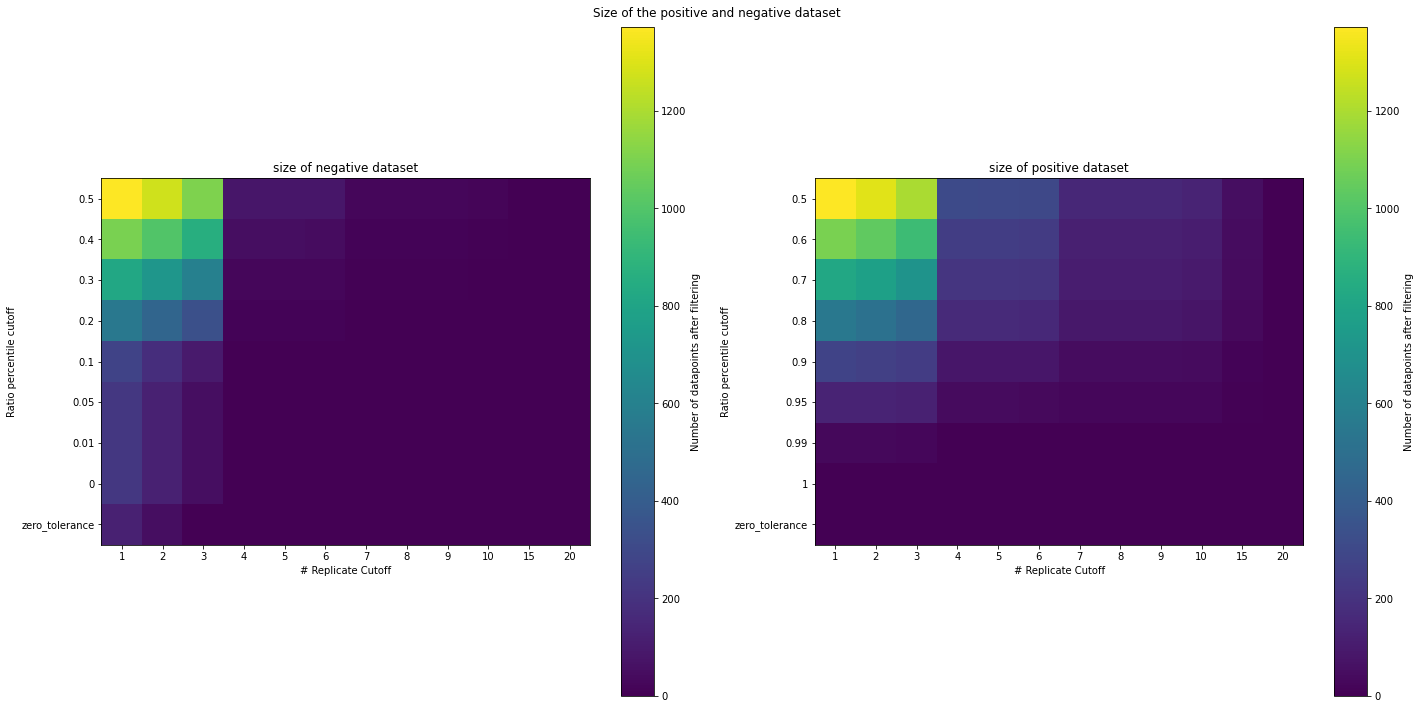

**Figure S4: Dataset hyperparameters for Adams et al. (2016) eLife.**

**Adams 2016**

**Figure S5: Dataset hyperparameters for Jenson et al. (2018) PNAS.**

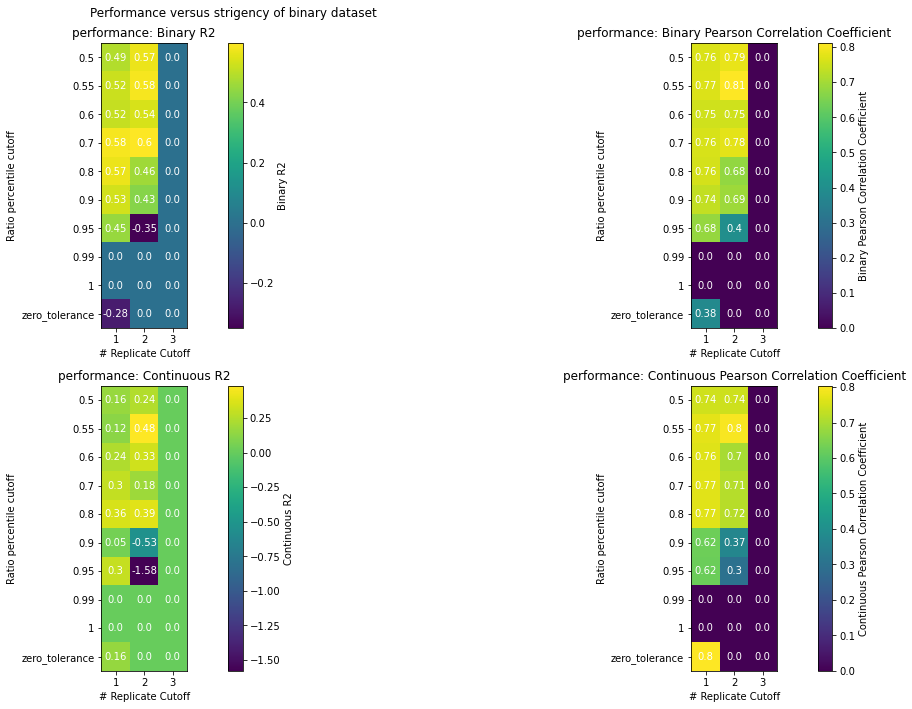

**Jenson 2018**

**Bfl-1**

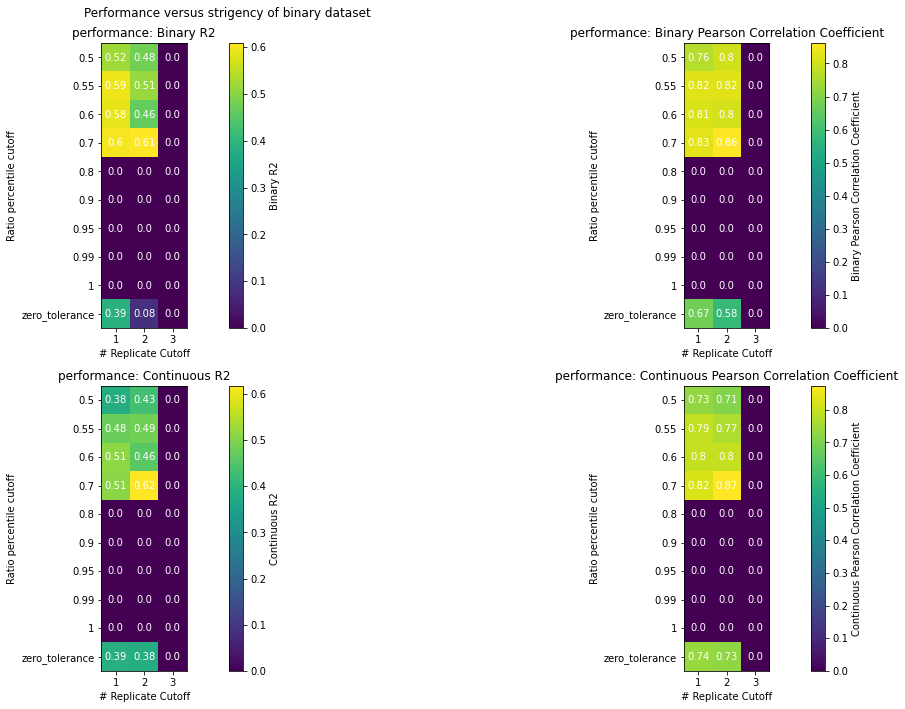

**Mcl-1**

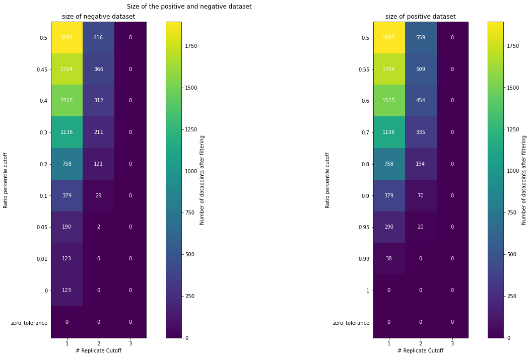

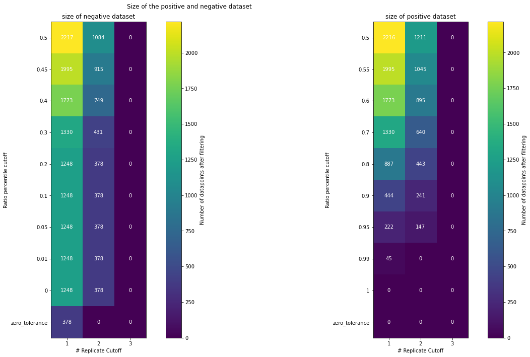

**Bcl-xL**

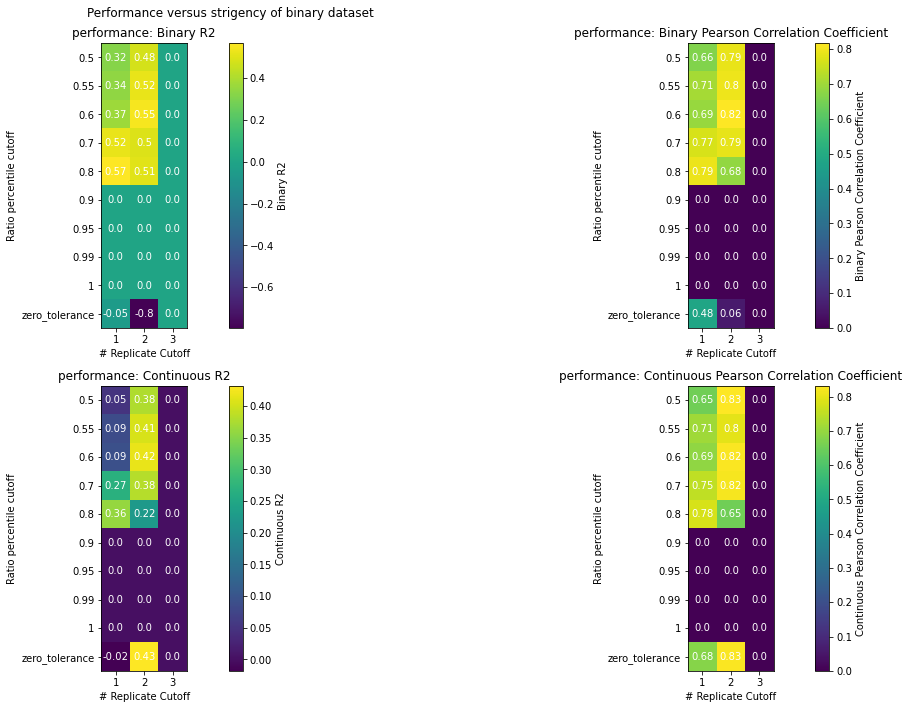

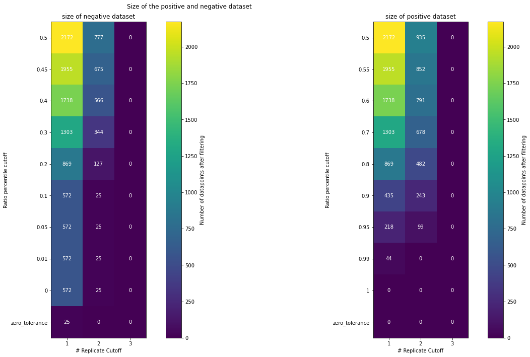

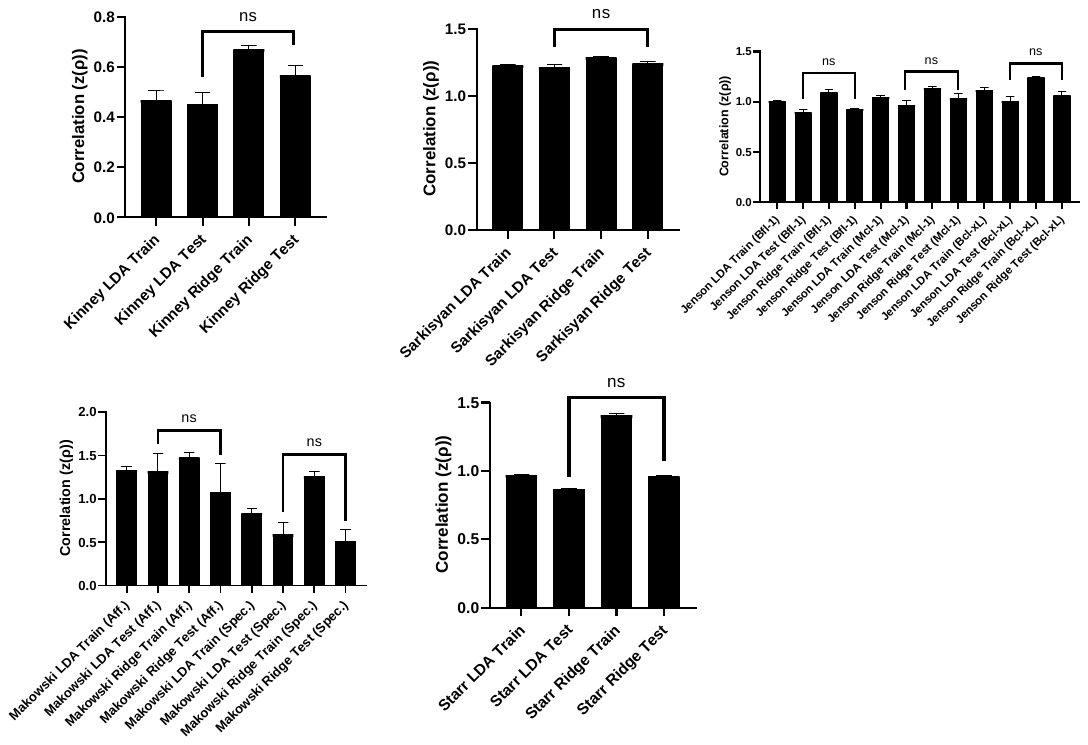

**Figure S6: Training and Test Set Performance statistics**

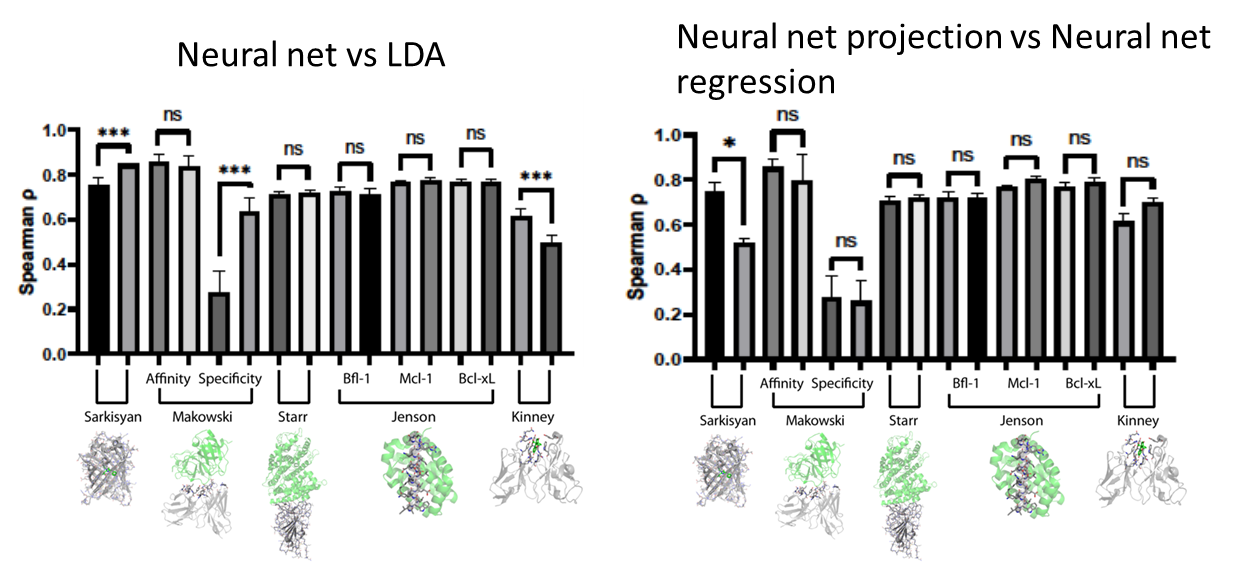

**Figure S7: Neural net performance statistics. (Left)** Neural net performance statistics versus linear discriminant analysis for test set. For all datasets, neural nets are on the left and LDA is on the right. **(Right)** Neural net classifier prediction of continuous mode versus neural net regressor for test set. For all datasets, neural net classifiers are on the left and the regressors are on the right. (*: p < 0.05, ***: p<0.0001).

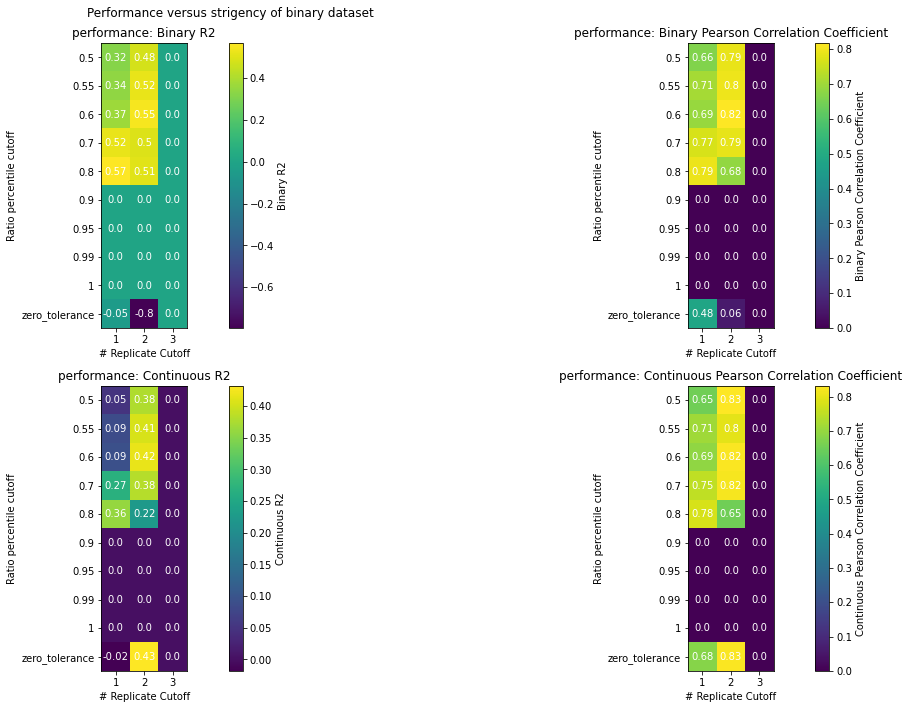

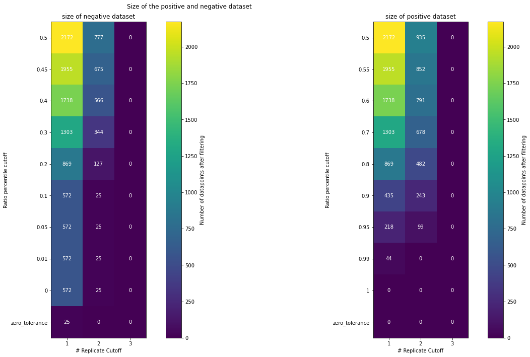

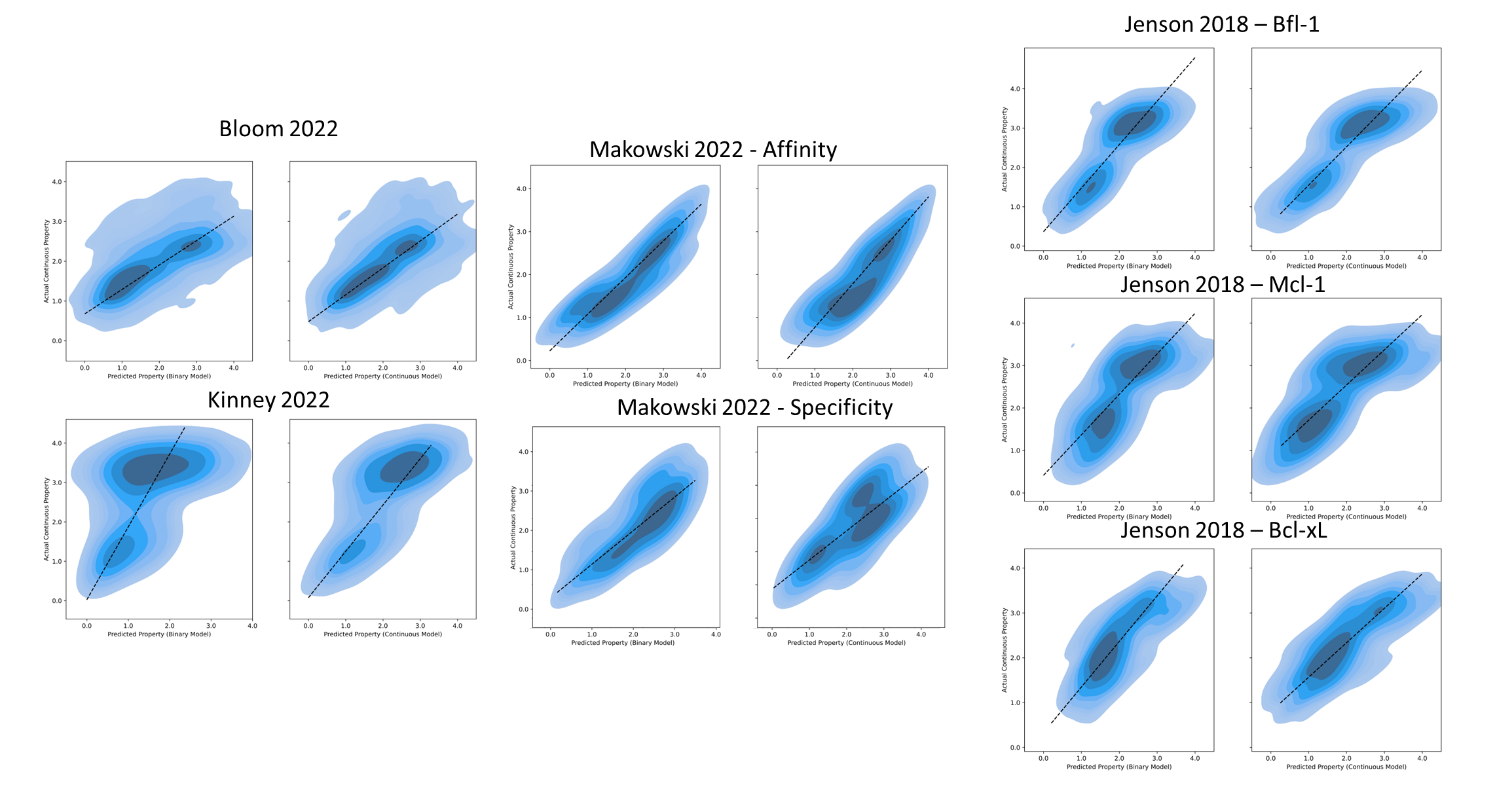

Bcl-xL

**Figure S8: Kernel density estimates for linear discriminant models projects’ correlations with continuous protein property values.**

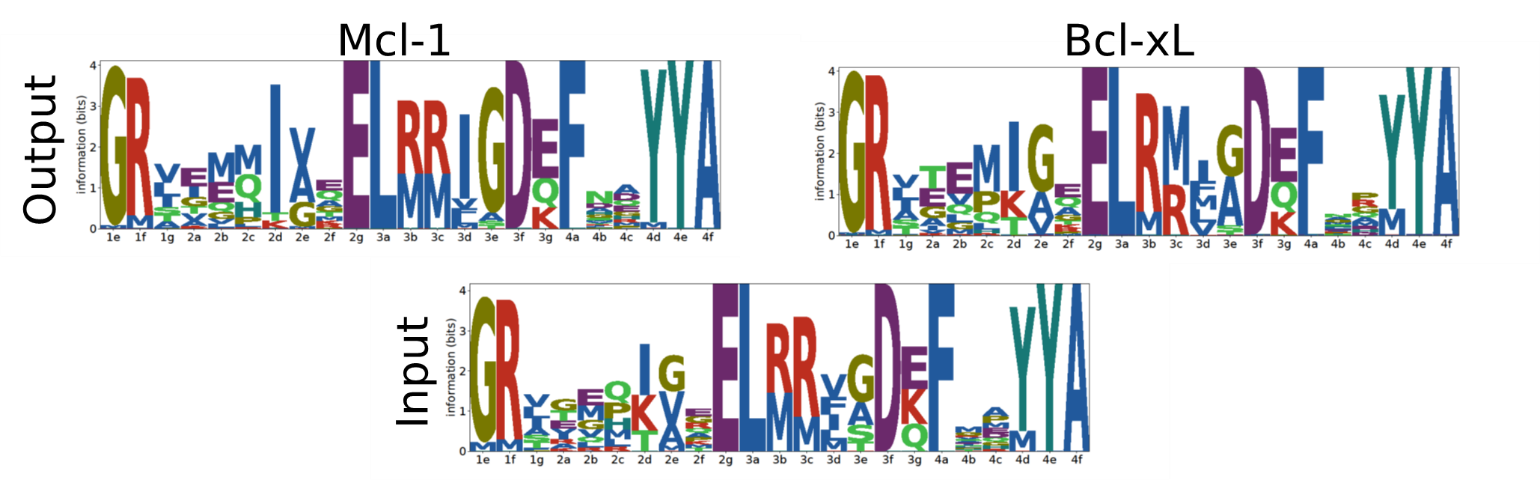

**Figure S9: Logoplots of input and output (pre- and post- sorting, respectively) for Bcl-xL and Mcl-1 stapled peptide libraries**

Mcl-1

Bcl-xL

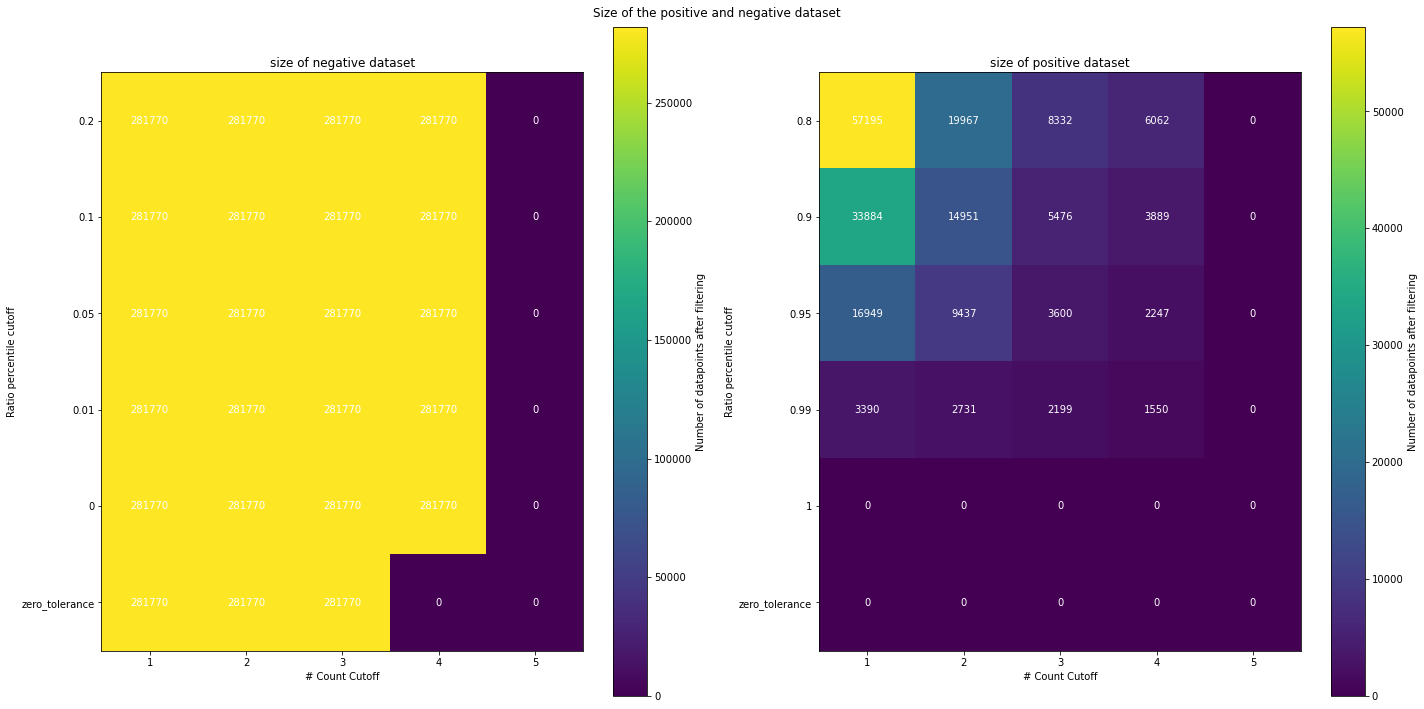

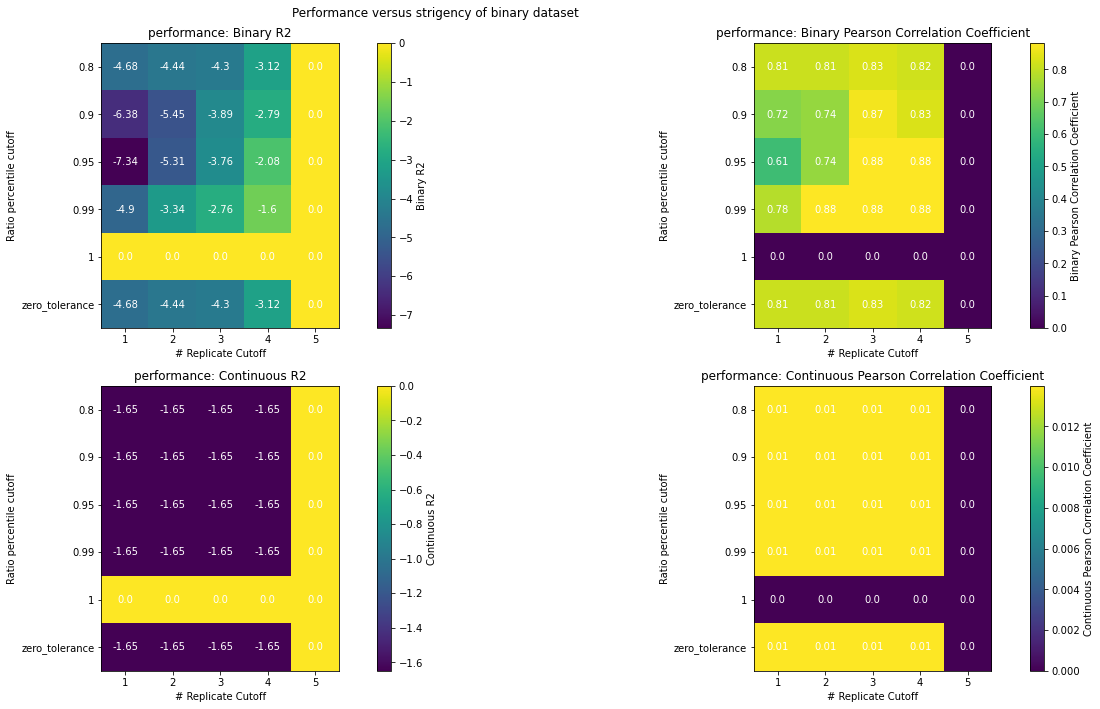

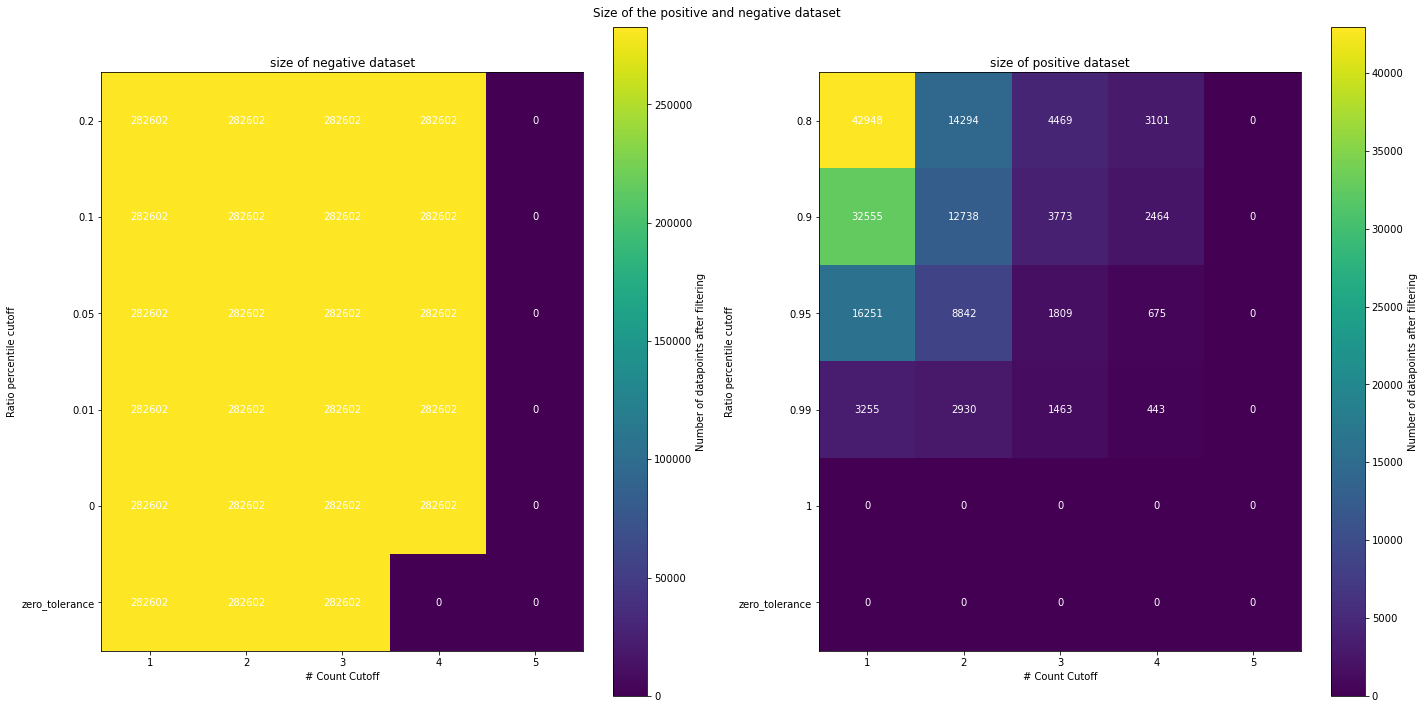

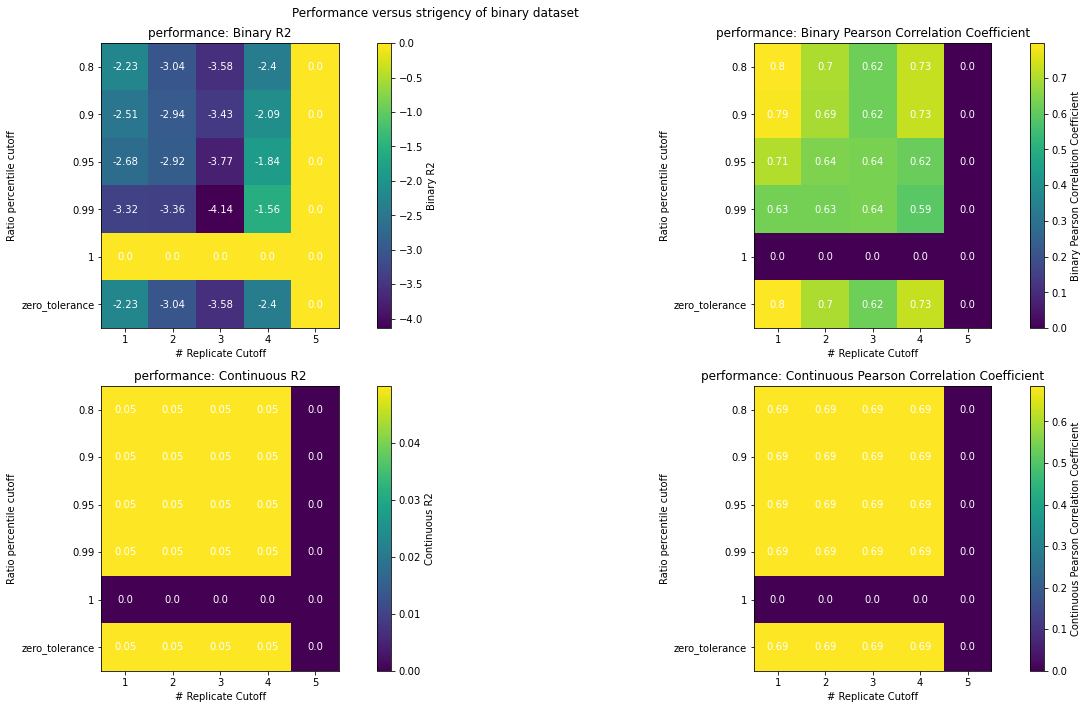

**Figure S10: Dataset hyperparameter data size and performance for pro-apoptotic anti-Bcl-2 stapled peptide libraries.**

**Figure S11: Random variants from Mcl-1 and Bcl-xL FACS 2-4 for low-throughput continuous binding measurement via bacterial cell surface and flow cytometry.**

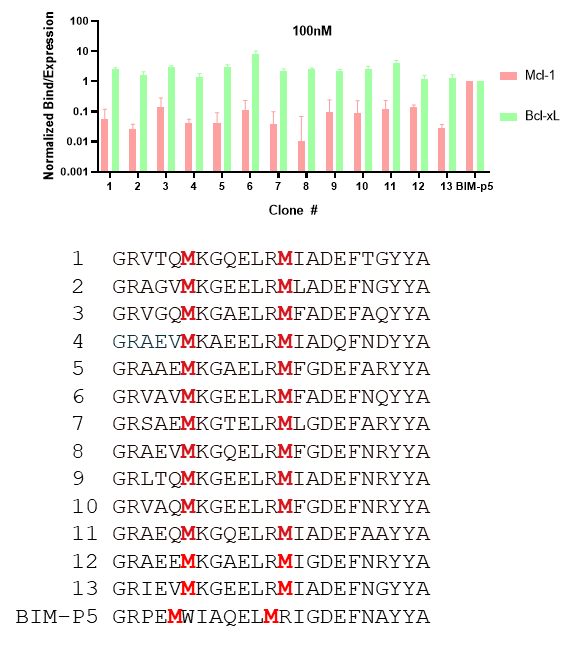

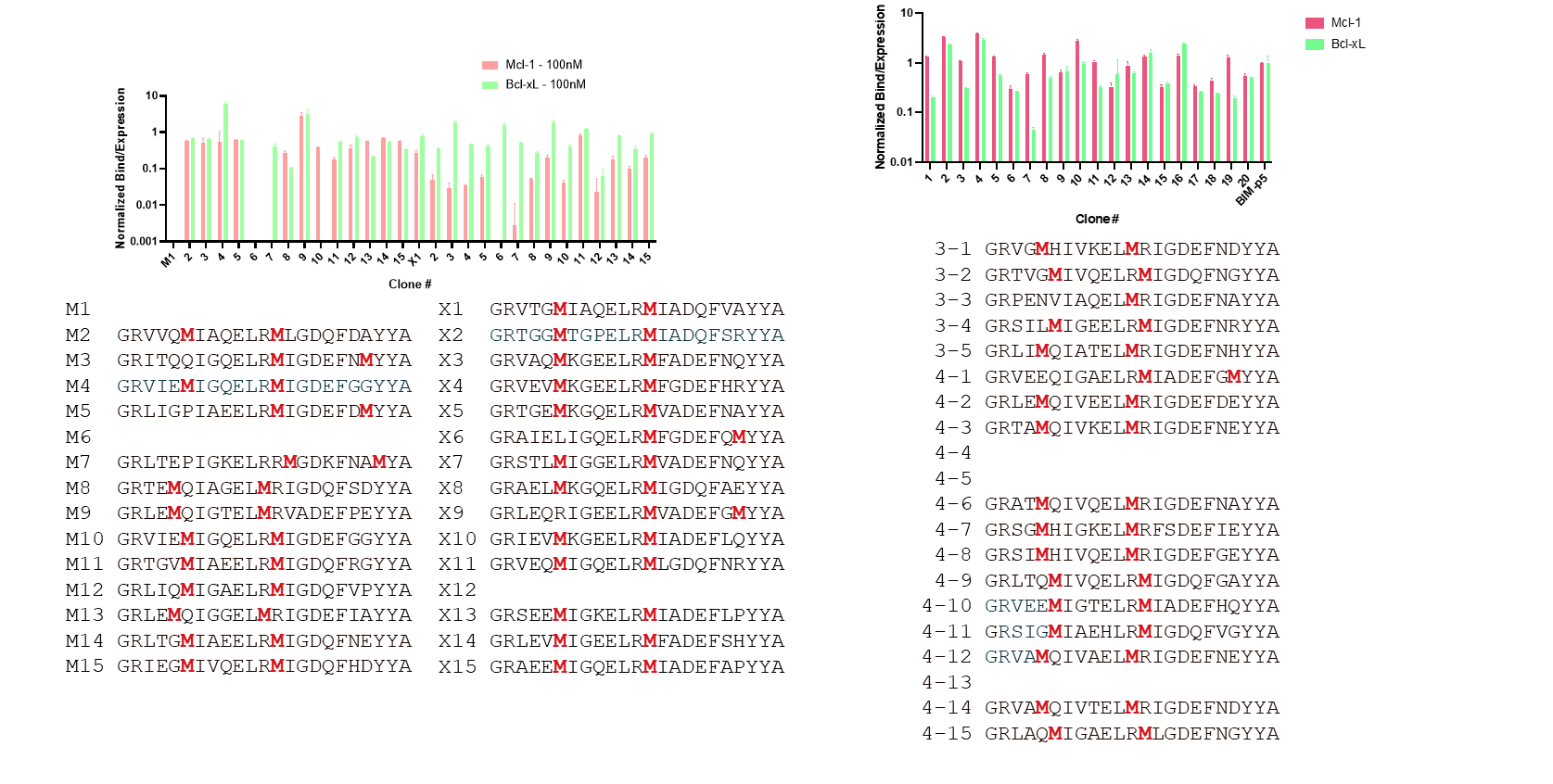

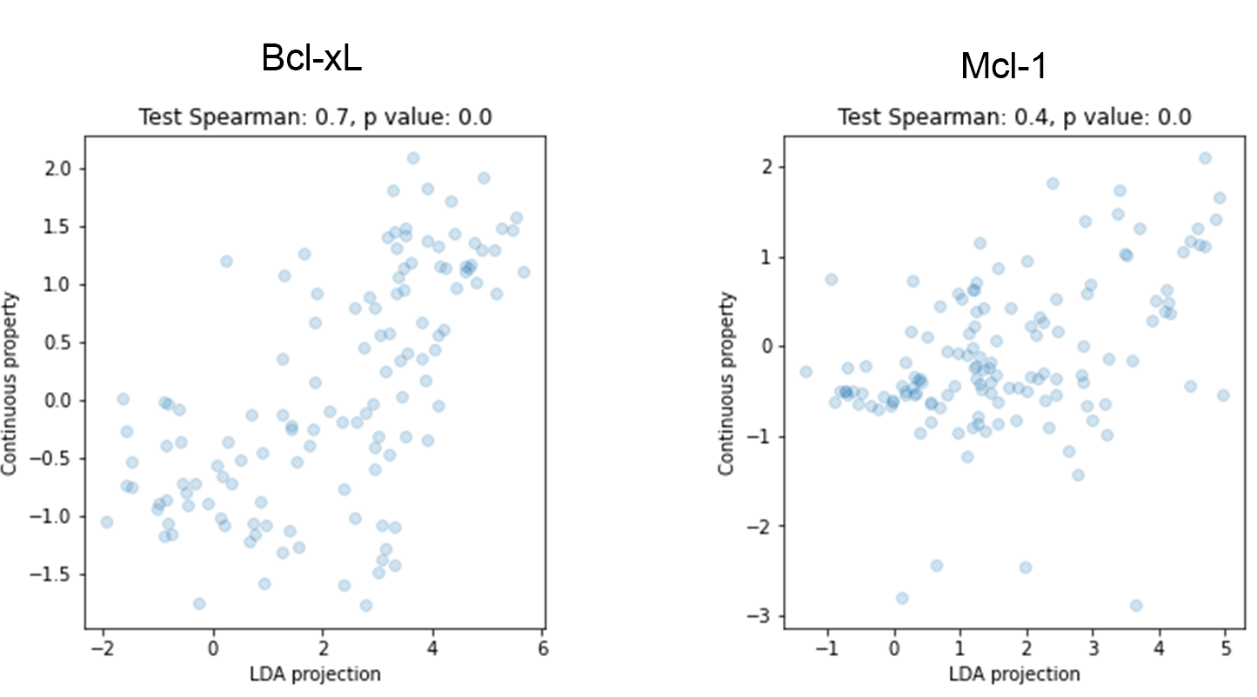

**Figure S13: LDA Projections from binary sorting versus multi-gate predicted continuous affinity (see Jenson et al. 2018 PNAS for more details).**

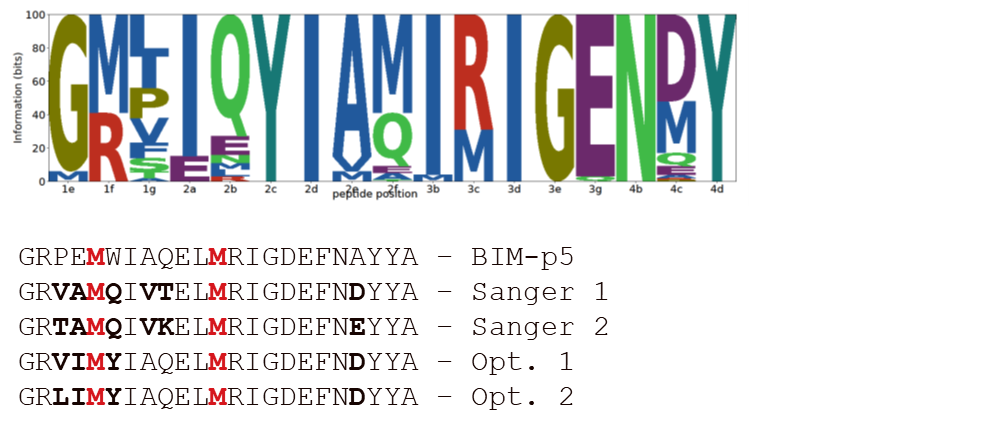

**Figure S14: LDA Projections from binary sorting versus multi-gate predicted continuous affinity (see Jenson et al. 2018 PNAS for more details).**

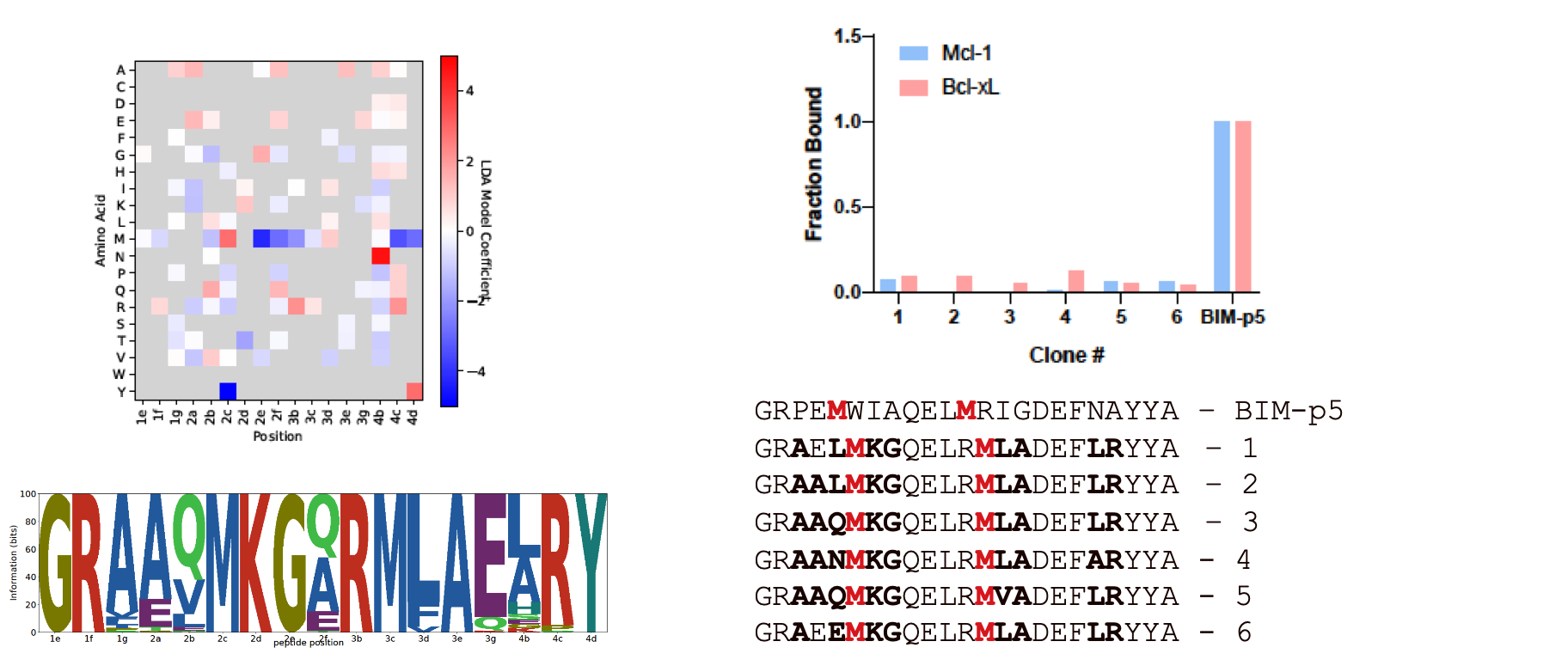

**Figure S15: Initial designs for Bcl-xL using ILP yielded non-binding sequences for both Bcl-xL and Mcl-1.** The optimization problem was set up as a maximization of Bcl-xL affinity subject to a low cutoff of Mcl-1 binding.

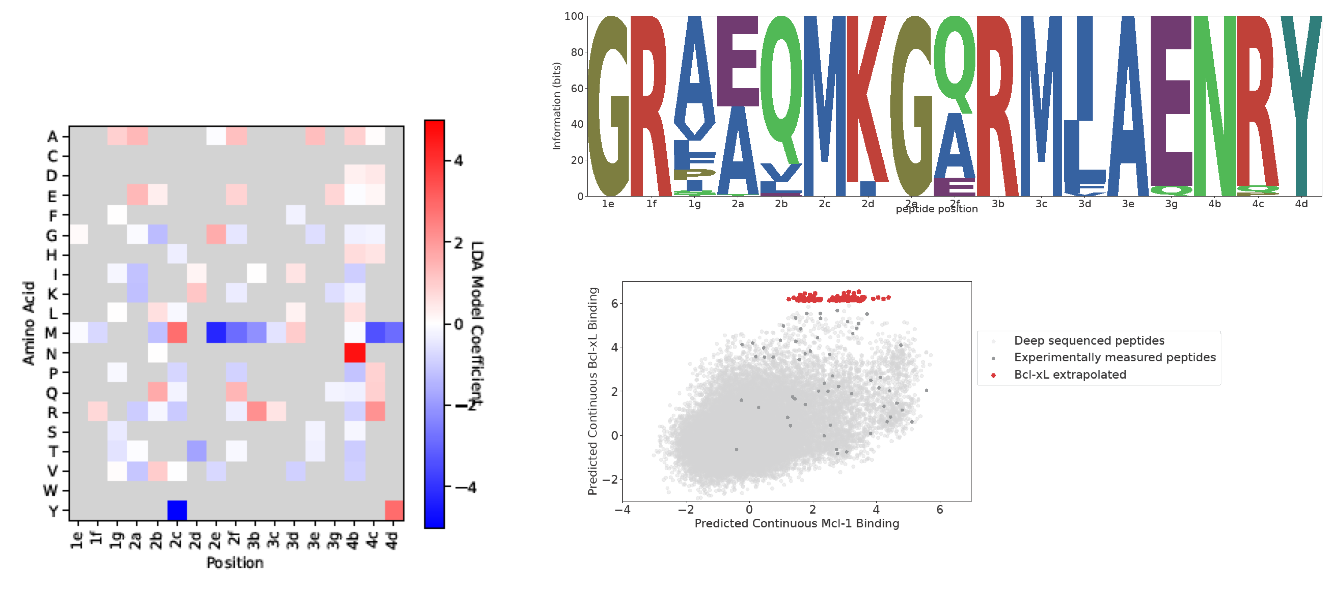

**Figure S16: Second iteration for Bcl-xL specific stapled peptides using ILP.**

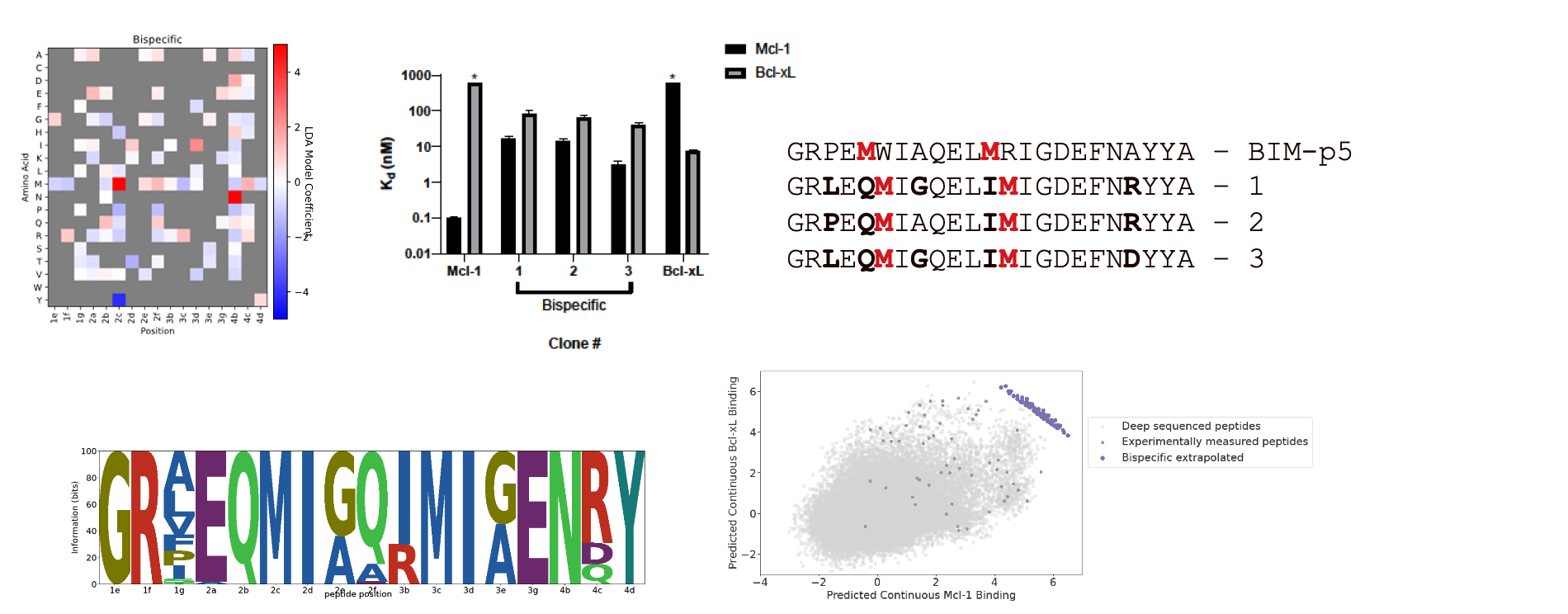

**Figure S17: Bispecific peptides designed via ILP.** The ILP objective was set as the maximization of both Mcl-1 and Bcl-xL score.

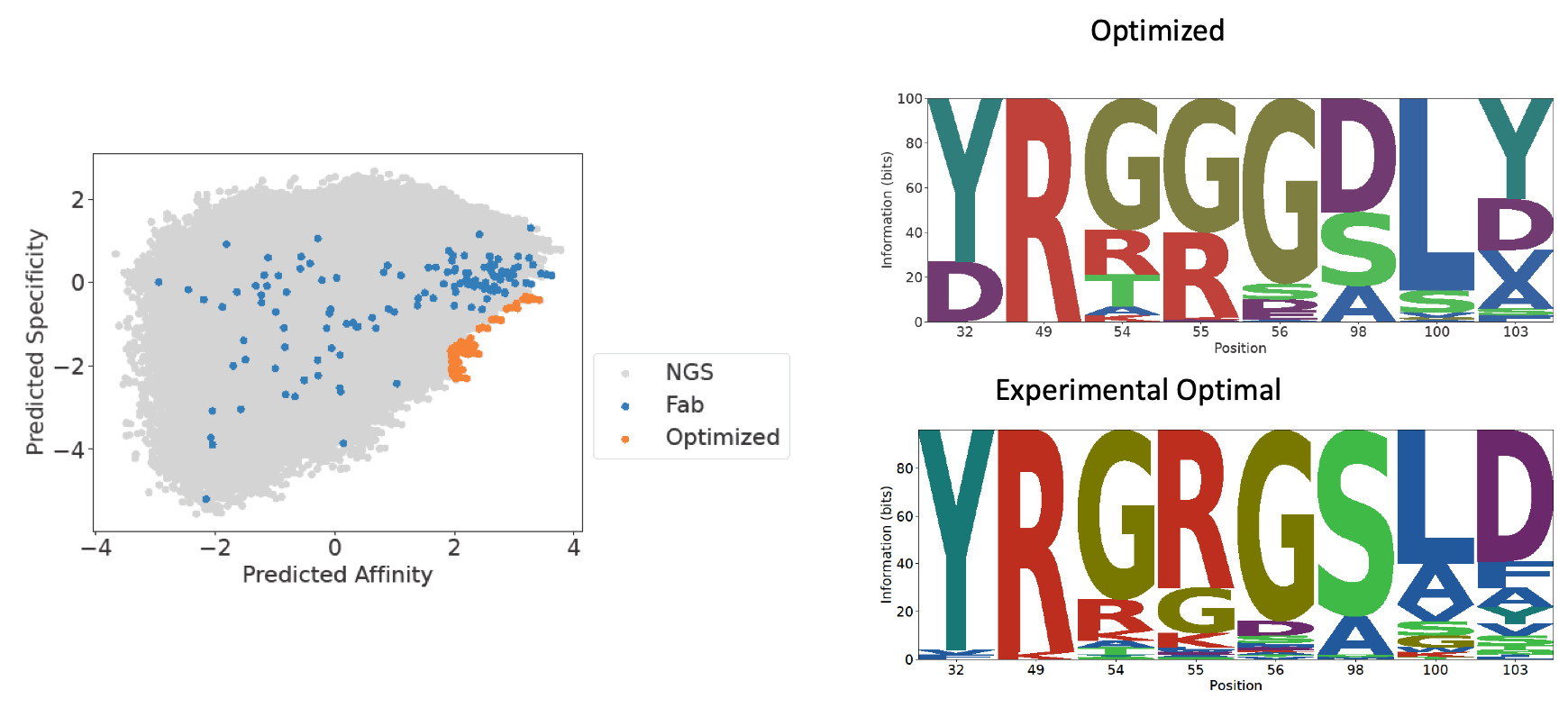

**Figure S17: High affinity and specificity antibodies from Makowski et al. (2022) Nature Communications via ILP.** The ILP objective was set as the maximization of affinity and minimization of specificity, subject to a minimum affinity threshold.
